# The cellular poly(rC)-binding protein 2 prevents the early steps of hepatitis C virus virion assembly

**DOI:** 10.1101/2022.04.12.488029

**Authors:** Sophie E. Cousineau, Selena M. Sagan

## Abstract

Poly(rC)-binding protein 2 (PCBP2) was previously shown to bind to the hepatitis C virus (HCV) genome; however, its precise role in the viral life cycle remained unclear. Herein, we found that PCBP2 does not directly affect viral entry, translation, genome stability, replication, or virion egress. Rather, our data suggests that endogenous PCBP2 normally limits virion assembly, thereby indirectly promoting translation and replication by increasing the translating/replicating pool of viral RNAs. Additionally, we found that an alternative RNA conformation (SLII^alt^) was important for efficient virion assembly, but functions in a PCBP2-independent manner. The latter may explain why the Japanese fulminant hepatitis 1 isolate is able to produce infectious particles in cell culture, while other HCV isolates are lost in translation. Taken together, our results suggest that PCBP2 and SLII^alt^ independently modulate HCV genome packaging and alter the balance of viral RNAs in the translating/replicating pool and those engaged in virion assembly.

**Graphical Abstract:** 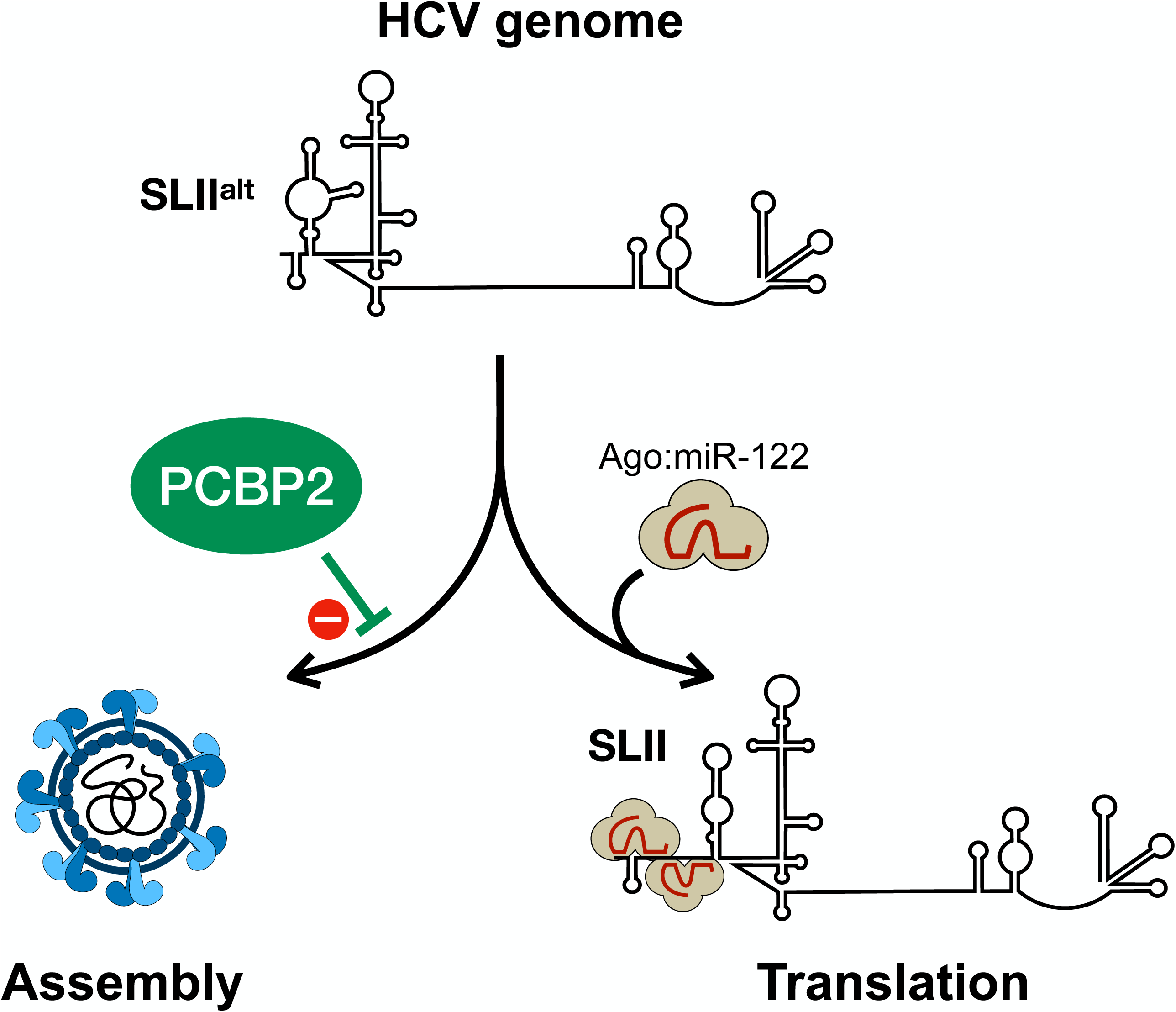

## INTRODUCTION

Hepatitis C virus (HCV) is an enveloped, positive-sense RNA virus of the *Flaviviridae* family (genus: hepacivirus) that typically causes a persistent liver infection (Simmonds et al., 2017). Its ∼9.6 kb genome contains a single open reading frame flanked by highly structured 5’ and 3’ untranslated regions (UTRs). The 5’ UTR contains stem-loop (SL) structures necessary for RNA replication (SLI-SLII) and translation (SLII-IV), while the 3’ UTR is implicated in viral replication and is composed of a hypervariable region, a poly(U/UC) tract, and a highly conserved 3’ X-tail. In the 5’ UTR, SLII-IV form an internal ribosomal entry site (IRES) that drives translation of the viral polyprotein, which is subsequently processed into 10 mature viral proteins, including 3 structural proteins (core, E1 and E2 glycoproteins), and 7 nonstructural proteins (p7, NS2, NS3, NS4A, NS4B, NS5A and NS5B) (Tsukiyama-Kohara et al., 1992; Lancaster et al., 2006). The structural proteins form the nucleocapsid and viral envelope, NS2 and NS3-4B are involved in polyprotein processing, while the NS3-5B proteins form the viral replicase, required for viral RNA replication (Lohmann et al., 1999). Additionally, the p7, NS2, NS3, NS4A and NS5A proteins have all been implicated in viral genome packaging and the assembly of progeny viral particles (Jones et al., 2007, 2011; Roder et al., 2019; Masaki et al., 2008; Appel et al., 2008)

Like all viruses, HCV is highly dependent on the host cell and co-opts numerous cellular proteins, RNAs and lipids to complete its life cycle. One important host factor is the liver-specific microRNA, miR-122, which binds to two sites in the 5’ UTR of the HCV genome (Jopling et al., 2005, 2008). These interactions promote HCV RNA accumulation, and miR-122 has been shown to have at least three roles in the viral life cycle: 1) it acts as a RNA chaperone (or riboswitch) to suppress an energetically stable RNA secondary structure (termed SLII^alt^) and allows the functional IRES (SLII-IV) to form; 2) it protects the 5’ terminus of the genome from pyrophosphatase activity and subsequent exoribonuclease-mediated decay; and 3) it promotes translation through interactions between the Argonaute (Ago):miR-122 complex at site 2 and the HCV IRES (Amador-Cañizares et al., 2018a, 2018b; Kincaid et al., 2018; Schult et al., 2018; Chahal et al., 2019; Henke et al., 2008; Roberts et al., 2011). Specific mutations in the 5’ UTR can circumvent some of these roles, by enabling the spontaneous formation of the functional SLII structure and/or by increasing base-pairing at the 5’ terminus to protect the viral genome from exoribonuclease-mediated decay, even in the absence of miR-122 (Chahal et al., 2021). In addition to these roles, miR-122 has been proposed to modulate viral RNA replication by facilitating the formation of replication organelles, potentially by facilitating the switch from translation to viral RNA replication, although the mechanism by which this occurs has not yet been fully elucidated (Panigrahi et al., 2022; Ono et al., 2017; Masaki et al., 2015; Li et al., 2019).

The poly(rC)-binding protein 2 (PCBP2) is one of the three most abundant cellular RNA-binding proteins with a strong affinity for poly(rC), along with its paralogs hnRNP K and PCBP1 (Matunis et al., 1992; Makeyev et al., 1999). PCBP2 is a multifunctional protein that can bind to hundreds of cellular RNAs, including its own mRNA transcript (Waggoner and Liebhaber, 2003). Depending on the context of its binding site and binding partners, PCBP2 can modulate the stability and/or translation of its mRNA targets (Wang et al., 1999; Gonzalez-Moro et al., 2020). Beyond its cellular mRNA interactions, PCBP2 has also been reported to interact with several viral RNAs, and has been demonstrated to participate in viral translation, RNA stability, replication and/or viral assembly (Beura et al., 2011; Palusa et al., 2012; López-Manríquez et al., 2013; Pingale et al., 2020; Zell et al., 2008; Lin et al., 2008; Graff et al., 1998; Collier et al., 1998; Woolaway et al., 2007). PCBP2’s role in the poliovirus (PV) life cycle is particularly well-characterized, where it mediates the switch from translation to replication (Gamarnik and Andino, 1997; Parsley et al., 1997; Blyn et al., 1997; Bedard et al., 2004; Perera et al., 2007; Chase et al., 2014). Specifically, PCBP2 is both an essential IRES trans-acting factor and part of the complex that circularizes the genome to initiate viral RNA replication, and viral protease-mediated PCBP2 cleavage triggers the switch from translation to replication. The importance of PCBP2 in the PV life cycle, and its interactions with several other viral RNAs, prompted investigations into whether PCBP2 played a similar role in the HCV life cycle.

In a previous siRNA screen, PCBP2 knockdown was found to initially enhance HCV titers and viral RNA levels, followed by a sharp decline in viral RNA accumulation and infectious particle production (Randall et al., 2007). However, the precise mechanism by which PCBP2 modulates HCV RNA accumulation and viral particle production remained elusive, as studies examining viral translation and RNA replication using different experimental systems arrived at contradictory conclusions (Rosenfeld and Racaniello, 2005; Fukushi et al., 2001; Wang et al., 2011; Shirasaki et al., 2010; Choi et al., 2004; Fontanes et al., 2009; Masaki et al., 2015). Furthermore, while miR-122’s promotion of viral RNA accumulation was reported to rely on endogenous PCBP2, the relationship between PCBP2 and miR-122, and whether PCBP2’s role in the HCV life cycle depends on miR-122, also remains unclear (Masaki et al., 2015). While PCBP2 has been known to bind the HCV 5’ UTR for more than two decades, its precise binding sites were only recently mapped by cross-linking immunoprecipitation (iCLIP), which identified six conserved PCBP2 binding sites across two HCV genotypes (Spångberg and Schwartz, 1999; Flynn et al., 2015). Specifically, two binding sites were identified in the 5’ UTR (near SLI, and over the initiation codon in SLIV of the IRES), three sites were identified in the polyprotein-coding region (in the *core*, *E2* and *NS5B* genes), and one site is located in the 3’ UTR (overlapping the variable region and poly(U/UC)-tract) (Flynn et al., 2015). However, despite the identification of these PCBP2 interaction sites, the role of PCBP2 in the HCV life cycle remains unclear. Thus, herein we aimed to clarify the role of PCBP2 in the HCV life cycle.

Using HCVcc, we found that endogenous PCBP2 was important for optimal HCV RNA accumulation and infectious particle production in cell culture. By examining individual steps of the viral life cycle, we ruled out a direct role for PCBP2 in viral entry, translation, genome stability, viral RNA replication, or virion secretion. Rather, we discovered that only viral RNAs that undergo the early steps of viral packaging were sensitive to PCBP2 knockdown, suggesting that endogenous PCBP2 normally limits HCV assembly, and thus indirectly increases viral translation and viral RNA accumulation by increasing the translating/replicating pool of viral RNAs. In addition, in an effort to better understand PCBP2’s interactions with the HCV genome at the 5’ terminus, we inadvertently revealed that a previously identified alternative conformation (SLII_alt_) was required for efficient HCV assembly. However, the importance of SLII^alt^ in virion assembly appears to act upstream or function in a PCBP2-independent manner. Thus, we propose a new model for HCV virion assembly that suggests that SLII^alt^ and PCBP2 both modulate HCV genome packaging, and thereby alter the balance between viral RNAs in the translating/replicating pool and those engaged in virion assembly.

## MATERIALS AND METHODS

### Cell culture

Huh-7.5 human hepatoma cells were provided by Dr. Charlie Rice (Rockefeller University) and maintained in complete media: Dulbecco’s Modified Eagle Media (DMEM) supplemented with heat-inactivated 10% fetal bovine serum (FBS), 2 mM L-glutamine, and 1X MEM non-essential amino acids. Human embryonic kidney cells (293T) were provided by Dr. Martin J Richer (McGill University) and maintained in DMEM supplemented with 10% FBS. All cells were maintained at 37°C/5% CO_2_ and were routinely screened for mycoplasma contamination.

### Plasmids and viral RNAs

The pJFH-1_T_ plasmid encodes a cell culture-adapted Japanese Fulminant Hepatitis (JFH-1; HCV genotype 2a) with three adaptive mutations that increase viral titers in cell culture (Russell et al., 2008). The pJ6/JFH1 FL RLuc WT (“RLuc-wt”) and pJ6/JFH-1 FL RLuc GNN (“RLuc-GNN”) plasmids bear full-length viral sequences derived from the J6 (structural genes) and JFH-1 (nonstructural genes) isolates of HCV, with a *Renilla* luciferase (RLuc) reporter inserted between the p7 and NS2-coding regions. RLuc-GNN also bears and inactivating GNN mutation within the NS5B RNA polymerase active site (Jones et al., 2007). The pJ6/JFH-1 mono RLuc-NS2 (“ Δcore-p7”) and pJ6/JFH-1 E1-p7 del (“ΔE1-p7”) plasmids – truncated versions of the *Renilla* reporter virus with deletions of structural genes through p7 – were provided by Dr. Joyce Wilson (University of Saskatchewan, Saskatoon, SK, Canada) (Panigrahi et al., 2022). The pJ6/JFH Δcore (“ΔCore”) plasmid consists of a truncated version of the *Renilla* reporter virus with a deletion of the core-coding gene that retained the first 15 codons (necessary for a functional HCV IRES) and the final 14 codons (which orient the E1 protein in the ER) of the core-coding sequence. To clone ΔCore, the *EcoRI* to *KpnI* fragment was subcloned to a temporary plasmid and PCR amplified with Q5 high fidelity DNA polymerase (NEB) using the dCore-BamHI-FW (5’-TTT CTG GAT CCT TGC TGG CCC TGC TGT CCT GCA TC-3’) and dCore-BamHI-RV (5’-GGG CGG GAT CCG GTG TTT CTT TTG GTT TTT CTT TGA G-3’) primers, and was auto-ligated after *BamHI* digestion. The *EcoRI* to *KpnI* fragment was then subcloned back into the parental pJ6/JFH-RLuc plasmid. The pJ6/JFH-1 FL RLuc-NS5A-GFP (“NS5A-GFP”) plasmid, a full-length *Renilla* reporter virus with a GFP insertion within the NS5A domain III, was subcloned as previously described (Cousineau et al., 2022).

The pJFH1T-U4C, pJFH1T-G20A and pJFH1T-G28A plasmids encode full-length JFH-1_T_ genomes with either the U4C, G20A, or G28A mutation. To introduce the U4C mutation, the pJFH-1_T_ plasmid was amplified using Q5 high fidelity DNA polymerase (NEB) and the EcoRI- T7-JFH1-U4C-FW (5’- GGC CAG TGA ATT CTA ATA CGA CTC ACT ATA GAC CCG CCC C-3’) and JFH-IRES-REV (5’- CGC CCT ATC AGG CAG TAC CAC AA-3’) primers, then digested with *EcoRI* and *AgeI* and ligated back into the pJFH-1_T_ plasmid. A similar procedure was followed to introduce the G20A mutation, with the modification that the pJFH-1_T_ template was first amplified with the JFH1-G20A-FW primer (5’- CAC TAT AGA CCT GCC CCT AAT AGG GGC AAC ACT -3’) and JFH-IRES-REV, followed with another amplification with the EcoRI-T7-FW (5’- GGC CAG TGA ATT CTA ATA CGA CTC ACT ATA GAC CTG CCC C-3’) and JFH-IRES-REV primers prior to digestion and ligation. To make the G28A mutant, the J6/JFH-RLuc-G28A plasmid (which already contained the G28A mutation within a JFH-1 5’ UTR (Chahal et al., 2021)) was amplified with the EcoRI-T7-JFH-FW and JFH-IRES-REV primers, and the product was digested and ligated into pJFH-1_T_.

To make uncapped viral RNAs, all plasmid templates were linearized with *XbaI* and were *in vitro* transcribed with T7 RNA polymerase (NEB). Briefly, 1 µg of linear template DNA was incubated at 30°C for 1 h with 200 U T7 RNA polymerase, 1 mM each of ATP, CTP, and UTP, 1.2 mM GTP, and 50 U RiboLock RNAse inhibitor (ThermoFisher Scientific), followed by a 15 min DNaseI (NEB) digestion at 37°C. The firefly luciferase (FLuc) mRNA was transcribed from the Luciferase T7 Control DNA plasmid (Promega) linearized using *XmnI* and *in vitro* transcribed using the mMessage mMachine T7 Kit (Life Technologies) according to the manufacturer’s instructions.

### Generation of infectious HCV stocks

To generate viral stocks, 30 µg of *in vitro* transcribed JFH-1_T_ RNA was transfected into Huh-7.5 cells using the DMRIE-C reagent (Life Technologies) according to the manufacturer’s instructions. Four days post-transfection, infectious cell supernatants were passed through a 0.45 µm filter and infectious viral titers were determined by focus-forming unit assay (Russell et al., 2008). Infectious virus was amplified for two passages through Huh-7.5 cells at a multiplicity of infection (MOI) of 0.1. Viral stocks were aliquoted and stored at -80°C until use.

### Focus-forming unit (FFU) assays

One day prior to infection, 8-well chamber slides (Lab-Tek) were seeded with 4 x 10^5^ Huh-7.5 cells/well. Infections were performed with 10-fold serial dilutions of viral samples in 100 µL for 4 h, after which the supernatant was replaced with fresh media. Three days post-infection, slides were fixed in 100% acetone and stained with anti-HCV core antibody (1:100, clone B2, Anogen), and subsequently with the AlexaFluor-488-conjugated anti-mouse antibody (1:200, ThermoFisher Scientific) for immunofluorescence analysis. Viral titers are expressed as the number of focus-forming units (FFU) per mL.

Extracellular virus titers were determined directly from cell supernatants, while intracellular virus titers were determined after cell pellets were subjected to lysis via four freeze-thaw cycles, removal of cellular debris by centrifugation and recovery of virus-containing supernatants.

### MicroRNAs and siRNA sequences

siGL3 (siCTRL): 5’-CUU ACG CUG AGU ACU UCG AUU-3’, siGL3* : 5’-UCG AAG UAC UCA GCG UAA GUU-3’, miR122_p2-8_ (siCTRL for luciferase experiments): 5’-UAA UCA CAG ACA AUG GUG UUU GU-3’, miR122_p2-8_*: 5’-AAA CGC CAU UAU CUG UGA GGA UA-3’ (Machlin et al., 2011), siPCBP2-1: 5’-UCC CUU UCU GCU GUU CAC CUU-3’, siPCBP2-1*: 5’-GGU GAA CAG CAG AAA GGG AUU-3’, siPCBP2-2: 5’-GGA CAG UAU GCC AUU CCA CUU-3’, and siPCBP2-2*: 5’-GUG GAA UGG CAU ACU GUC CUU-3’ (Randall et al., 2007) were all synthesized by Integrated DNA Technologies.

All microRNA and siRNA duplexes were diluted to a final concentration of 20 µM in RNA annealing buffer (150 mM HEPES pH 7.4, 500 mM potassium acetate, 10 mM magnesium acetate), annealed at 37°C for 1 h and stored at -20°C. Annealed siPCBP2-1 and siPCBP2-2 were mixed together at a 1:1 ratio prior to transfection. For all knockdown experiments, 50 nM siRNA transfections were conducted 2 days prior to infection or electroporation of viral RNAs. Transfections were conducted using the Lipofectamine RNAiMAX (Invitrogen) according to the manufacturer’s instructions with the modification that 20 µL of reagent were used to transfect a 10-cm dish of cells.

### HCV and VSV pseudoparticles (HCVpp and VSVpp)

HCVpp consisting of a Firefly luciferase reporter lentiviral vector pseudotyped with the HCV E1 and E2 glycoprotein (from the H77, a genotype 1a strain) were a kind gift from Dr. John Law (University of Alberta) (Hsu et al., 2003). To generate lentiviral vectors pseudotyped with the VSV-G glycoprotein (VSVpp), a 90% confluent 10-cm dish of 293T cells were transfected with 10 µg pPRIME-FLuc, 5 µg psPAX.2, and 2.5 µg pVSV-G plasmid with 10 µL Lipofectamine 2000 (Invitrogen) diluted in 4 mL Opti-MEM. Media was changed 4, 20, and 28 h post-transfection. At 48 h post-transfection, the cell culture media was passed through a 0.45 µm filter and stored at -80°C.

To assay for cell entry, HCVpp and VSVpp were diluted 1/3 in dilution media (1X DMEM, 3% FBS, 100 IU penicillin and 100 µg/mL streptomycin) with 20 mM HEPES and 4 µg/µL polybrene, and then introduced to Huh-7.5 cells by spinoculation at 1,200 rpm for 1 h at room temperature. The cells were left to recover at 37°C for at least 5 h before the pseudoparticle-containing media was changed for fresh complete Huh-7.5 media. In parallel, cells seeded in a 6-well plate were transfected with 1 µg of pPRIME-FLuc plasmid using Lipofectamine 2000 (Invitrogen) according to the manufacturer’s instructions. Three days post-spinoculation and transfection, cells were lysed in passive lysis buffer (Promega) and firefly luciferase activity was assayed using the Dual Reporter Luciferase kit (Promega).

### Infections

Three days prior to infection, 10-cm dishes were seeded with 5 x 10^5^ Huh-7.5 cells, which were transfected with siRNA duplexes on the following day. On the day of infection, each 10-cm dish – containing approximately 1 x 10^6^ cells – was infected with 5 x 10^4^ FFU of JFH-1_T_ diluted in 3 mL complete media. Four to five hours post-infection, each infected plate was split into three 10-cm dishes. Protein, RNA, and virus samples were collected three days post-infection.

### Electroporations

For each electroporation, 6 x 10^6^ cells resuspended in 400-600 µL cold PBS were mixed with 5 µg of replication-competent viral RNA, or with 10 µg of nonreplicative GNN J6/JFH-1 RLuc RNA with 2 µg of FLuc mRNA, and electroporated in 4-mm cuvettes at 270 V, 950 µF, infinite resistance optimized for the Bio-Rad GenePulser XCell (Bio-Rad). Electroporated cells were resuspended in complete Huh-7.5 media and transferred to 6-well plates for luciferase assays and protein expression analyses, or to a 10-cm dish to assess infectious virus production.

### Inhibition of RNA replication by 2’CMA

Two days post-siRNA transfection, Huh-7.5 cells were infected with JFH-1_T_ at a MOI of 0.05. Four to five hours post-infection, each plate of infected cells was split into 6-well plates. Three days post-infection, the media on these cells was changed for complete Huh-7.5 media with 5 µM 2’CMA (2’C-methyladenosine, Carbosynth), an HCV NS5B polymerase inhibitor, or DMSO (vehicle control) (Carroll et al., 2003). Total RNA and intracellular virus samples were collected at 0, 6 and 12 h post-treatment, while cell culture supernatants were collected 6 and 12 h post-treatment. Protein samples were collected from untreated plates to assess PCBP1 knockdown efficiency by Western blot.

### RNA structure predictions

To predict the secondary structures and free energy (ΔG) of the first 117 nucleotides of the JFH-1 sequence, the RNAstructure 6.4 secondary structure prediction software was accessed from the Matthews lab server https://rna.urmc.rochester.edu/RNAstructureWeb/ (Bellaousov et al., 2013). To avoid spurious interactions that artificially decreased the calculated ΔG, the first three nucleotides of the sequence were constrained to be single-stranded. The results were saved as dot bracket files and used to generate the predicted structures with the RNA2Drawer browser app https://rna2drawer.app/ (Johnson et al., 2019).

### Western blot analysis

To collect total intracellular protein samples, cells were lysed in RIPA buffer (150 mM sodium chloride, 1% NP-40, 0.5% sodium deoxycholate, 0.1% SDS, 50 mM Tris pH 8.0) supplemented with Complete Protease Inhibitor Cocktail (Roche) and frozen at -80°C. Cellular debris was pelleted by centrifugation at 16,000 x g for 30 min at 4°C, and the supernatant protein concentration was quantified by BCA Protein Assay (ThermoScientific). Ten micrograms of sample were separated on 10-12% SDS-PAGE gels prior to transfer onto Immobilon-P PVDF membranes (Millipore). Membranes were blocked in 5% milk and incubated overnight with primary antibodies diluted in 5% BSA: mouse anti-PCBP2 (clone 5F12, Abnova H00005094-M07, diluted 1:20,000); rabbit anti-actin (A2066, Sigma, 1:20,000); mouse anti-HCV core (clone B2, Anogen MO-I40015B, 1:7,500); mouse anti-JFH-1 NS5A (clone 7B5, BioFront Technologies, 1:10,000). Blots were incubated for 1 h with HRP-conjugated secondary antibodies diluted in 5% milk: anti-mouse (HAF007, R&D Systems, 1:25,000); anti-rabbit (111-035-144, Jackson ImmunoResearch Laboratories, 1:50,000) and visualized using enhanced chemiluminescence (ECL Prime Western Blotting Detection Reagent, Fisher Scientific).

### RNA isolation and Northern blot analysis

Total RNA was harvested using TRIzol Reagent (ThermoFisher Scientific) according to the manufacturer’s instructions. Ten micrograms of total RNA were separated on a 1% agarose gel containing 1X 3-(N-morpholino)propanesulfonic acid (MOPS) buffer and 2.2 M formaldehyde and transferred to a Zeta-probe membrane (Bio-Rad) by capillary transfer in 20X SSC buffer (3 M NaCl, 0.3 M sodium citrate). Membranes were hybridized in ExpressHyb Hybridization Buffer (ClonTech) to random-primed ^32^P-labeled DNA probes (RadPrime DNA labelling system, Life Technologies) complementary to HCV (nt 84-374) and γ-actin (nt 685-1171). Autoradiograph band densities were quantified using Fiji (ImageJ2 2.3.0) (Schindelin et al., 2012).

### RT-qPCR analysis

The iTaq Universal Probes One-Step kit (Bio-Rad) was used to perform duplex assays probing for the HCV genome (NS5B-FW primer: 5’-AGA CAC TCC CCT ATC AAT TCA TGG C-3’; NS5B-RV primer: 5’-GCG TCA AGC CCG TGT AAC C-3’; NS5B-FAM probe: 5’-ATG GGT TCG CAT GGT CCT AAT GAC ACA C-3’) (Cousineau et al., 2022) and the GAPDH loading control (PrimePCR Probe assay with HEX probe, Bio-Rad). Each 20 µL reaction contained 500 ng of total RNA, 1.5 µL of the HCV primers and probe, and 0.5 µL of the GAPDH primers and probe. RT-PCR reactions were conducted in a CFX96 Touch Deep Well Real-Time PCR system (Bio-Rad). Genome copies were calculated against a genomic RNA standard curve, and fold-differences in gene expression were calculated using the 2^−ΔΔCt^ method (Livak and Schmittgen, 2001).

### Luciferase assays

For translation and replication assays, cells were washed in PBS and harvested in 100 µL of 1X passive lysis buffer (Promega). The Dual-Luciferase Assay Reporter Kit (Promega) was used to measure both *Renilla* and firefly luciferase activity according to the manufacturer’s instructions with the modification that 25 µL of reagent were used with 10 µL of sample. All samples were measured in triplicate.

### Data analysis

Statistical analyses were performed using GraphPad Prism v9 (GraphPad, USA). Statistical significance was determined by paired t-test to compare results obtained from multiple experiments, and by two-way ANOVA with Geisser-Greenhouse and Bonferroni corrections when more than two comparisons were applied at once. Half-lives were calculated with a one-step decay curve calculated using a least-squares regression, and error was reported as the asymmetrical (profile likelihood) 95% confidence interval of the half-life. To calculate virion accumulation or secretion rates, a simple linear regression using the least squares method was performed; the resulting slope and standard error (in viral titers over time) represents the rate of virus accumulation/secretion.

## RESULTS

### PCBP2 is required for optimal HCVcc accumulation in cell culture

A previous siRNA screen identified PCBP2 as an important cellular factor for optimal HCV replication in cell culture (Randall et al., 2007). To further investigate the role of PCBP2, we knocked down endogenous PCBP2 in Huh-7.5 cells, infected with the JFH-1_T_ strain (HCVcc), and assessed viral protein expression, viral RNA accumulation, and extracellular viral titers (**Figure 1)**. Notably, transfecting cells with the anti-PCBP2 siRNA generally led to a transient increase in PCBP2 protein levels at day 1 post-transfection, followed by a sustained knockdown from days 2-5 post-transfection (**Figure S1**). As such, all HCV infections were carried out at day 2 post-siRNA transfection. Endogenous PCBP2 knockdown resulted in an approximately 2.3-fold decrease in viral protein and RNA accumulation in Huh-7.5 cells (**Figure 1A-D**). Consistent with this reduction in viral protein and RNA accumulation, we also observed a 1.7-fold decrease in viral titers (**Figure 1E**). Thus, in line with previous findings, we found that endogenous PCBP2 is necessary for optimal HCV replication in cell culture, although the precise step(s) of the viral life cycle modulated by PCBP2 remained unclear.

**Figure 1.**
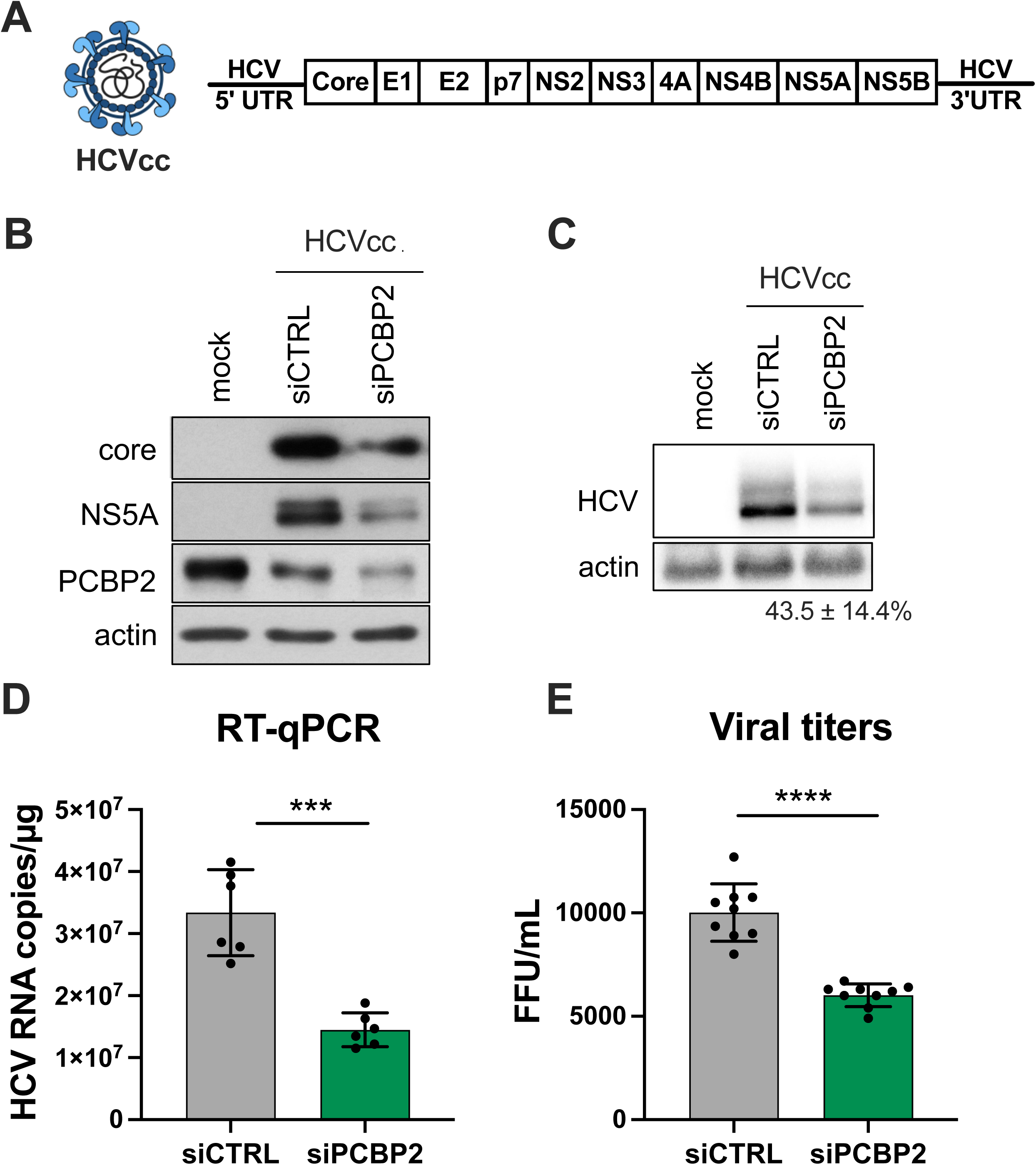
PCBP2 is required for optimal RNA accumulation and infectious virus production in cell culture. **(A)** Schematic representation of the HCVcc (JFH-1_T_) infectious particles and genomic RNA used in infections. Huh-7.5 cells were transfected with siPCBP2 or siCTRL 2 days prior to infection with JFH-1_T_ (MOI = 0.05). Viral protein, total RNA and extracellular infectious virus were harvested 3 days post-infection. **(B)** Viral protein expression analysis by Western blot. **(C)** Viral RNA accumulation analysis by Northern blot, and **(D)** quantification by RT-qPCR. **(E)** Extracellular (secreted) virus titers, quantified by FFU assay. All data are representative of three independent replicates, and error bars represent the standard deviation of the mean. P-values were calculated by paired t-test (*** p < 0.001; **** p < 0.0001).

### PCBP2 knockdown has no effect on HCV entry, translation, or genome stability

To clarify PCBP2’s role in the HCV life cycle, we used assays to specifically examine each step of the viral life cycle. Firstly, we explored if PCBP2 knockdown had any effect on virus entry using the HCV pseudoparticle (HCVpp) system, which consists of lentiviral vectors with a firefly luciferase reporter gene pseudotyped with the HCV E1 and E2 glycoproteins (Hsu et al., 2003). HCVpp enters cells by clathrin-mediated endocytosis after engaging with HCV-specific entry receptors; as such, we used vesicular stomatitis virus (VSV) pseudoparticles (VSVpp) as a control for effects on clathrin-mediated endocytosis. Additionally, to verify that PCBP2 knockdown did not affect luciferase reporter gene expression, we assessed firefly luciferase expression from cells directly transfected with a FLuc reporter plasmid. In all cases, we found that PCBP2 knockdown had no impact on luciferase activity (**Figure 2A**). This suggests that endogenous PCBP2 does not affect FLuc reporter gene expression, clathrin-mediated endocytosis, or HCVpp entry.

**Figure 2.**
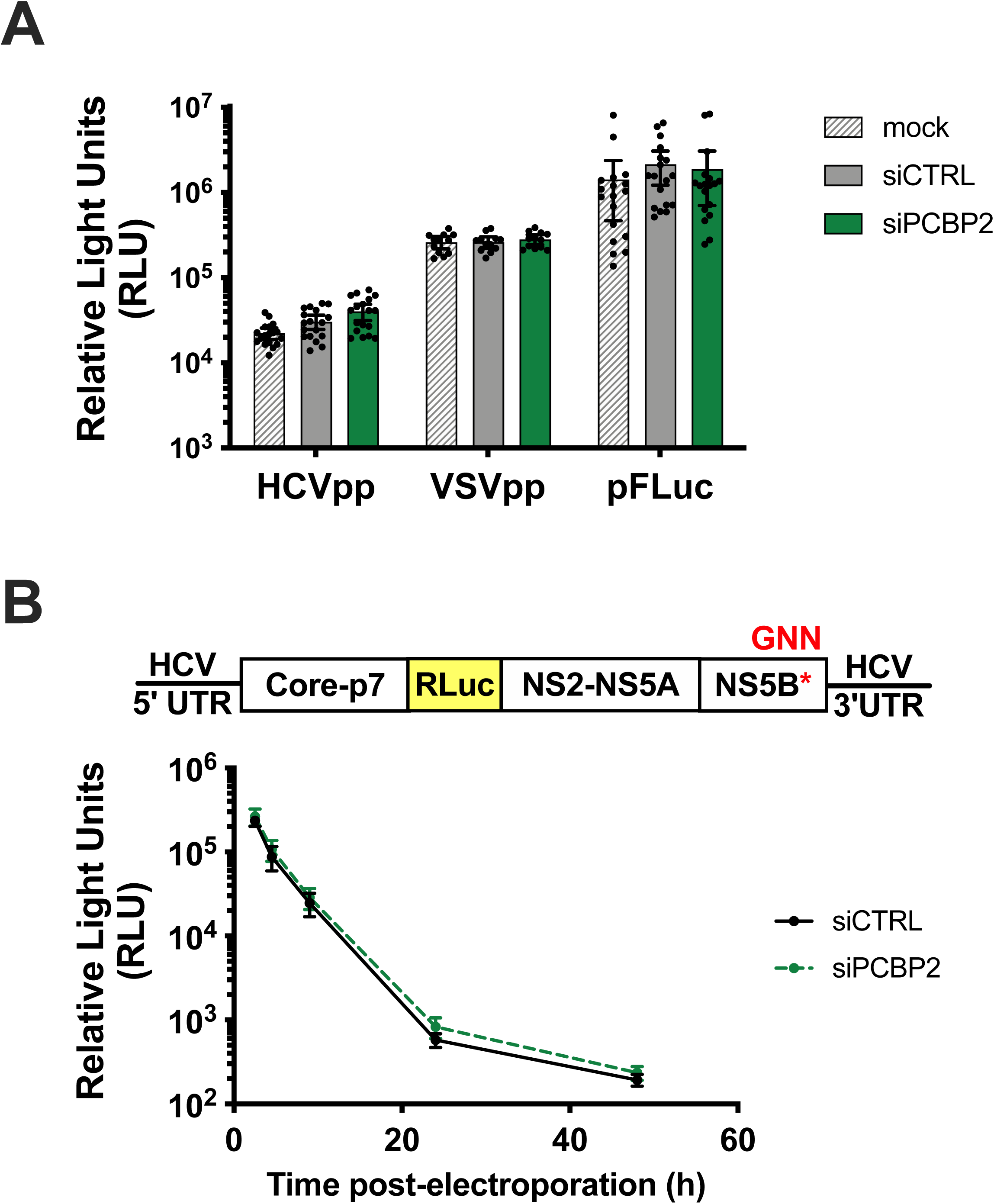
PCBP2 knockdown has no effect on HCV entry, translation, or genome stability. **(A)** Two days post-siRNA transfection, cells were spinoculated with luciferase reporter pseudoparticles bearing the HCV E1/E2 glycoproteins (HCVpp) or the VSV-G glycoprotein (VSVpp). In parallel, cells were transfected with a firefly luciferase expression plasmid. Samples were harvested 3 days post-infection/transfection, and analyzed by luciferase assay. The HCVpp and pFLuc data is representative of three independent replicates, while the VSVpp data is representative of two independent replicates. Error bars represent the standard deviation of the mean. **(B)** Huh-7.5 cells were electroporated with J6/JFH-RLuc-GNN reporter RNAs and a FLuc control mRNA, and luciferase activity was monitored at several timepoints post-electroporation. All data are representative of three independent replicates. Error bars represent the standard deviation of the mean. No statistically significant differences were found by paired t-test or two-way ANOVA.

PCBP2 is a known IRES trans-acting factor for several cellular and viral IRESes (Evans et al., 2003; Graff et al., 1998; Kim et al., 2018; Walter et al., 1999). However, it is still unclear whether PCBP2 affects HCV translation, as studies using a variety of experimental systems have arrived at contradictory conclusions (Choi et al., 2004; Fontanes et al., 2009; Fukushi et al., 2001; Masaki et al., 2015; Rosenfeld and Racaniello, 2005; Shirasaki et al., 2010; Wang et al., 2011). Thus, to specifically assess PCBP2’s effect on viral translation, we used full-length J6/JFH-1 RLuc reporter RNAs with an inactivating mutation in the NS5B polymerase active site (GNN). In this system, RLuc activity serves as a direct measure of HCV IRES-mediated translation, and over time this signal also serves as a proxy for viral RNA stability. We found that the siPCBP2 and siCTRL conditions had similar RLuc activity at all timepoints (**Figure 2B**). Additionally, we also assessed HCV IRES-mediated translation in isolation using PV, encephalomyocarditis (EMCV) and HCV IRES-luciferase reporter RNAs, which further confirmed that PCBP2 knockdown had no effect on HCV IRES-mediated translation (**Figure S2**). Finally, we observed that the half-lives of the J6/JFH-1 RLuc reporter RNAs were nearly identical, with 1.59 h (95% CI 1.13–2.21) and 1.44 h (95% CI 1.11–1.85) for the siPCBP2 and siCTRL conditions, respectively, suggesting that PCBP2 knockdown does not impact viral RNA stability. Thus, taken together, these results suggest that PCBP2 does not modulate either HCV IRES-mediated translation or viral genome stability.

### PCBP2 knockdown only affects the accumulation of packaging-competent viral RNAs

PCBP2 has been reported to bind to and promote the replication of viral RNAs through interactions with their 5’ UTRs (Walter et al., 2002; López-Manríquez et al., 2013). Since we had found that endogenous PCBP2 was important for HCV RNA accumulation during infection, but that it did not affect HCV IRES-mediated translation or genome stability (see **Figure 1C-D** and **Figure 2B**), we next examined if PCBP2 knockdown could influence viral RNA replication. To this end, we used the RLuc activity of replication-competent reporter RNAs as a proxy measure for viral RNA accumulation, starting with the full-length WT J6/JFH-1 RLuc RNA (**Figure 3A**). Consistent with our HCVcc infection experiments (**Figure 1**), we found that PCBP2 knockdown reduced luciferase expression of the full-length WT J6/JFH-1 RLuc viral RNA, with a >2-fold decrease in luciferase activity from days 1-3 post-electroporation (**Figure 3A**). Notably, this full-length reporter RNA is assembly competent, although the addition of the large RLuc gene severely impairs viral packaging efficiency (Cousineau et al., 2022; Herker et al., 2010; Poenisch et al., 2015). Thus, to examine viral RNA replication more specifically in the absence of any viral packaging, we assessed the luciferase activity of a subgenomic J6/JFH-1 RLuc replicon RNA lacking all the structural protein genes (Δcore-p7). Interestingly, we found that PCBP2 knockdown had no impact on RLuc expression or viral RNA accumulation of the Δcore-p7 subgenomic replicon RNA (**Figure 3B and Figure S3**). We observed a similar insensitivity to PCBP2 knockdown with a bicistronic subgenomic replicon, which lacks the *core* through *NS2*-coding region and whose expression of the NS3 through NS5B proteins is driven by the encephalomyocarditis virus (EMCV) IRES (**Figure S4**). Collectively, these results suggest that PCBP2 does not have a direct effect on viral RNA replication.

**Figure 3.**
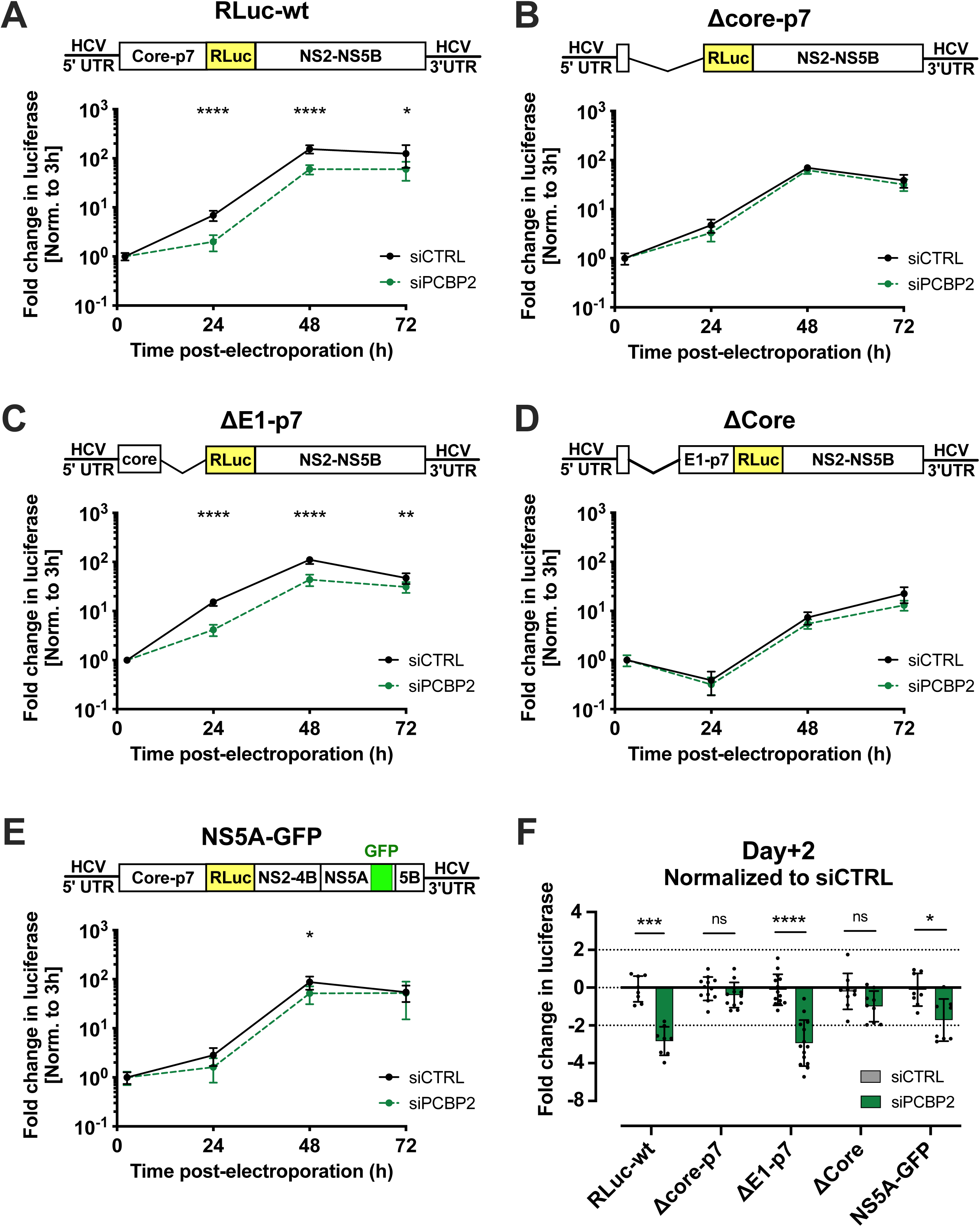
PCBP2 knockdown affects reporter viruses when the core-NS5A interaction is conserved. Two days post-siRNA transfection, Huh-7.5 cells were electroporated with **(A)** full-length RLuc WT J6/JFH-1 RNA, **(B)** Δcore-p7 J6/JFH-1 RLuc RNA, **(C)** ΔE1-p7 J6/JFH-1 RLuc RNA, **(D)** ΔCore J6/JFH-1 RLuc RNA, or **(E)** full-length NS5A-GFP J6/JFH-1 RLuc RNA. RLuc values were normalized to the early timepoint (3 h), to control for disparities in electroporation efficiency between experiments. **(F)** Comparison of luciferase values obtained 2 days post-electroporation, normalized to the siCTRL condition. Data are representative of three independent replicates; error bars represent the standard deviation of the mean. P-values were calculated by two-way ANOVA (ns, not significant; * p < 0.05; ** p < 0.01; *** p < 0.001; **** p < 0.0001).

At this point, it was unclear whether our results indicated that PCBP2’s effect on HCV RNA accumulation depended upon interactions with the core-p7 region of the viral RNA, or if its role in the viral life cycle was related to a specific effect on viral assembly and/or egress. Thus, to better understand why PCBP2 knockdown was detrimental to full-length viral RNA replication yet had no impact on the Δcore-p7 RNA, we further mapped the viral gene requirements for PCBP2 sensitivity, using subgenomic viral RNAs with deletions in E1 through p7 (ΔE1-p7) or the core-coding (ΔCore) region alone (**Figure 3C-D**). The ΔE1-p7 viral RNA was still sensitive to PCBP2 knockdown, with an approximately 2.5-fold decrease in luciferase activity and concomitant reduction in viral RNA accumulation (**Figure 3C and Figure S3**). In contrast, while ΔCore exhibited delayed replication kinetics, PCBP2 knockdown did not result in any further impairment of viral RNA replication (**Figure 3D**). Taken together, these results indicate that only viral RNAs that contain the *core* gene are sensitive to PCBP2 knockdown; however, it was still unclear if the *core*-coding sequence or the activity of the core protein was important for PCBP2 sensitivity.

Finally, we aimed to clarify if the *core* gene or if the packaging activity of the core protein was the relevant determinant of PCBP2 sensitivity. We reasoned that PCBP2 might regulate virion assembly since it had been reported to interact with the NS5A protein *in vitro*, and NS5A is implicated in virion assembly by delivering the viral RNA to the core protein for packaging (Wang et al., 2011; Masaki et al., 2008; Tellinghuisen et al., 2008). To test this hypothesis, we assessed the impact of PCBP2 knockdown on a packaging-deficient viral RNA that still retained the *core* gene: a full-length J6/JFH-1 RLuc NS5A-GFP reporter RNA, which bears a GFP insertion into the NS5A domain III previously shown to reduce virion assembly without impacting RNA replication (**Figure 3E**) (Masaki et al., 2008; Moradpour et al., 2004). We found that PCBP2 knockdown only modestly decreased luciferase activity of this NS5A-GFP reporter RNA, with a smaller than 2-fold decrease at the peak of replication (**Figure 3E and 3F**). Taken together, our results suggest that PCBP2 knockdown does not impair viral RNA replication directly, but rather that it is only able to modulate the accumulation of packaging-competent viral RNAs.

### PCBP2 knockdown promotes virion assembly, but does not affect virion egress

While our model suggested that PCBP2 plays a role in the early stages of virion assembly, our data thus far could not rule out an additional role in viral egress. Thus, to directly examine virion assembly and viral egress, we inhibited viral RNA replication using the nucleoside analog 2’ C-methyladenosine (2’CMA) and monitored intracellular and extracellular viral titers (**Figure 4**) (Carroll et al., 2003; Cousineau et al., 2022). Specifically, we performed PCBP2 knockdown and HCVcc infections as we had done previously, but at 3 days post-infection, we replaced the culture media with media containing 5 µM 2’CMA (or DMSO, as a vehicle control) and collected total RNA, intracellular and extracellular virions over the following 12 hours (**Figure 4A**). In agreement with our previous findings, we observed an overall reduction in viral RNA accumulation during PCBP2 knockdown (0 h timepoint, siCTRL vs. siPCBP2, **Figure 4B**). Importantly, the 2’CMA treatment efficiently blocked viral RNA accumulation, which continued to increase under the control (DMSO) condition (6-12 h timepoints, **Figure 4B**). When we assessed intracellular titers, we were surprised to see that in the DMSO (vehicle control) condition, despite starting with a lower level of intracellular virions, PCBP2 knockdown resulted in a similar level of intracellular virion accumulation to the siCTRL condition at the experimental endpoint (i.e. 0 and 12 h time points, **Figure 4C**). Moreover, we observed a shallower decline in intracellular titers in the 2’CMA treatment during PCBP2 knockdown (**Figure 4C**). This was reflected in the overall intracellular virion accumulation rate and supports our hypothesis that PCBP2 modulates virion assembly (**Figure 4D**). However, when we assessed extracellular titers, we did not observe any significant differences in the viral secretion rates, suggesting that PCBP2 does not modulate virion egress (**Figure 4E-F**). Taken together, our results suggest that PCBP2 knockdown promotes virion assembly, but does not have a significant impact on virion egress post-assembly. Collectively, our results suggest that endogenous PCBP2 limits the early steps of virion assembly, likely during viral RNA transfer from NS5A to the core protein for genome packaging (**Figure 4G**).

**Figure 4.**
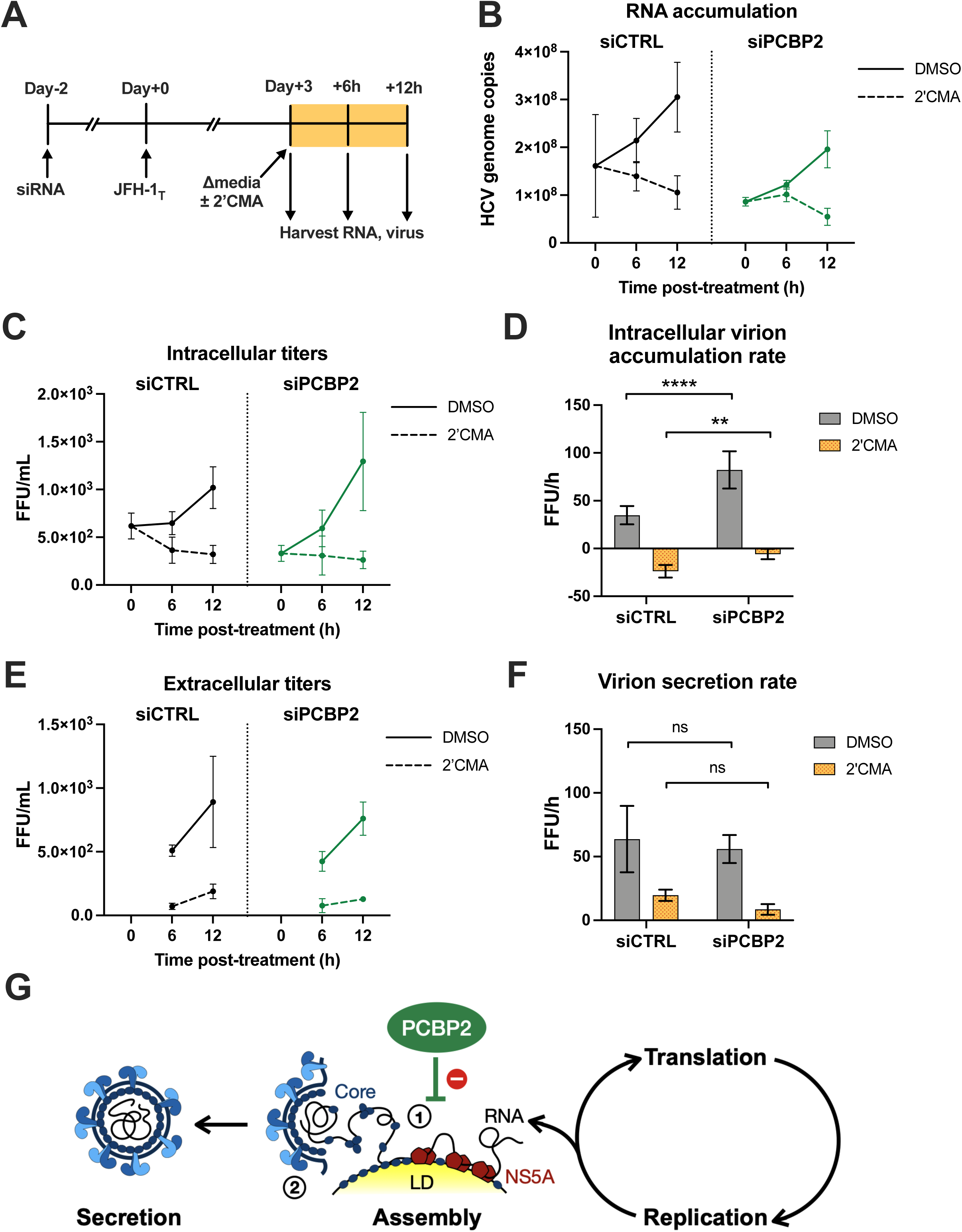
PCBP2 knockdown increases virion assembly, but does not affect virion egress. **(A)** Schematic representation of the experimental approach for 2’CMA experiments: two days post-siRNA transfection, Huh-7.5 cells were infected with JFH-1_T_ (MOI = 0.05). Three days post-infection, cells were treated with 2**’**CMA or DMSO (control). Total RNA, intracellular and extracellular virus was collected at t = 0, 6 and 12 h post-treatment. **(B)** Quantitative RT-PCR analysis after 2’CMA treatment. **(C)** Intracellular viral titers and **(D**) intracellular virus accumulation rate calculated by linear regression. **(E)** Extracellular viral titers and **(F)** virus secretion rates calculated by linear regression. **(G)** Model of PCBP2’s role in the HCV life cycle: PCBP2 normally inhibits the first stage of virion assembly, where the viral NS5A protein transfers the viral RNA to the core protein. All data are representative of three independent replicates and error bars in (B), (C) and (E) represent the standard deviation of the mean. Error bars in (D) and (F) represent the slopes of the linear regressions ± standard error. P-values were calculated by two-way ANOVA (ns, not significant; ** p < 0.01; **** p < 0.0001).

### The SLII^alt^ conformation promotes packaging into virions, independently of PCBP2’s role in virion assembly

Previous studies had mapped a PCBP2 binding site near the 5’ terminus of the HCV genome, a region of the viral RNA that contains two tandem miR-122 binding sites (Flynn et al., 2015; Jopling et al., 2008). Binding studies showed that PCBP2 and Ago:miR-122 competed to bind this stretch of the viral genome *in vitro*, with a more recent study clarifying that both Ago:miR-122 and PCBP2 could simultaneously occupy the 5’ terminus of the viral RNA (Masaki et al., 2015; Gebert et al., 2021). However, these binding studies used RNA probes that only contain nucleotides 1-47 of the HCV genome, which are unable to sample the alternative conformations that have previously been mapped to nucleotides 1-117 of the 5’ UTR (Chahal et al., 2019; Schult et al., 2018). In the current model for Ago:miR-122 interactions with the HCV genome, the first 117 nucleotides of the JFH-1 5’ UTR are predicted to take on one or more energetically stable conformation(s), collectively termed SLII^alt^, and Ago:miR-122 recruitment to the HCV genome stabilizes the SLII conformation, allowing the HCV IRES (SLII-IV) to form (Lagos-Quintana et al., 2002; Chahal et al., 2019; Schult et al., 2018) (**Figure 5A**). Importantly, we noticed that the putative PCBP2 binding site, a stretch of four cytidine residues (C41-44) is predicted to be single-stranded (accessible) in the SLII_alt_ conformations, but is partially base-paired, and thus inaccessible, in the canonical SLII conformation (**Figure 5A**, insets). Notably, nucleotides C41-45 are always predicted to be accessible in the 47 nucleotide long RNAs used in previous binding studies, as they are too short to form any alternate RNA structures (Masaki et al., 2015; Gebert et al., 2021). Thus, we were curious whether PCBP2’s effects on virion assembly were dependent upon the ability of the viral RNA to form the SLII^alt^ conformation, which exposes the putative PCBP2 binding site.

**Figure 5.**
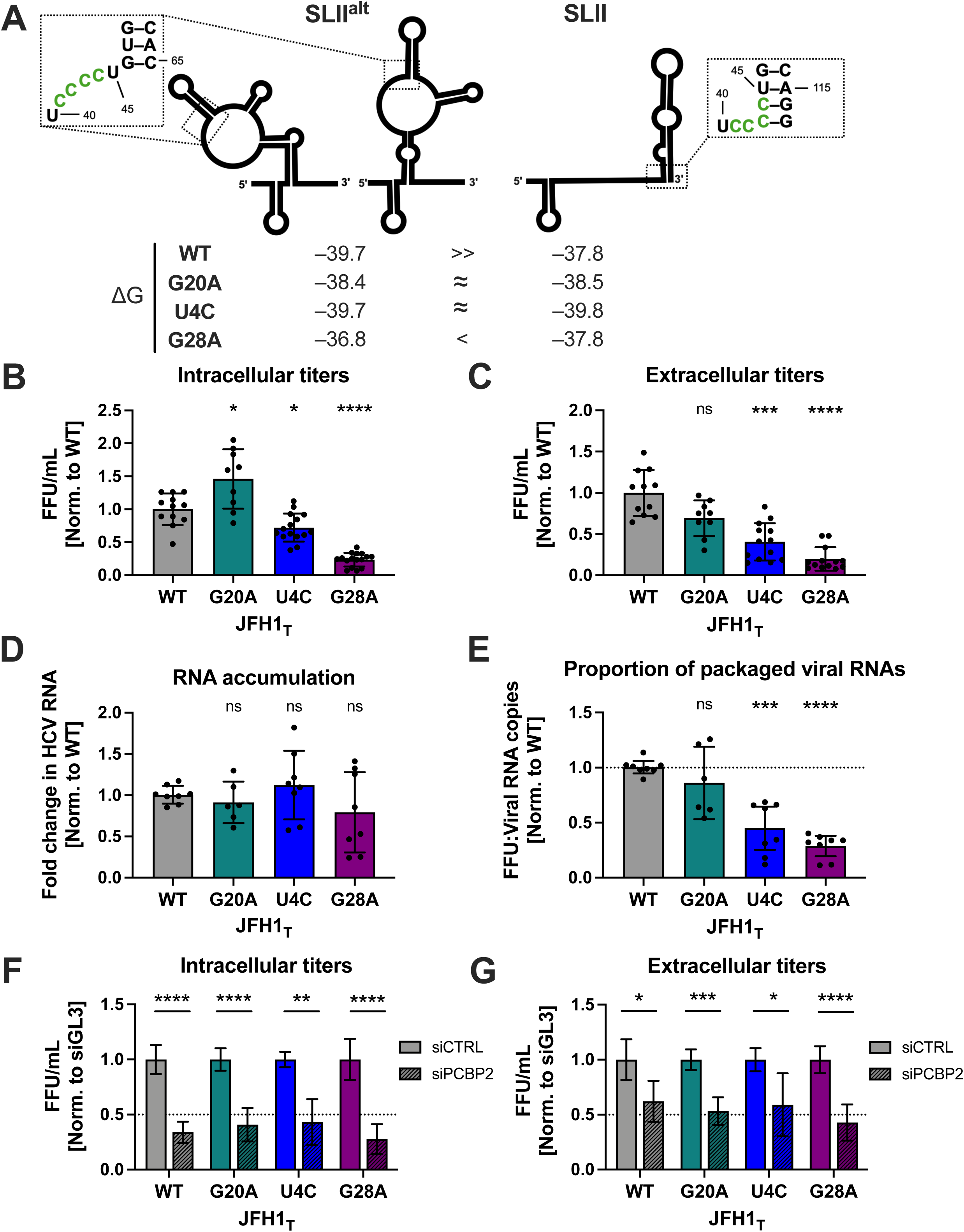
The SLII^alt^ conformation promotes virion assembly, but is not required for PCBP2’s effect on virion assembly. **(A)** RNA conformations of the first 117 nucleotides of the HCV 5**’** UTR, showing the free energies (ΔG) of SLII^alt^ and SLII for WT, G20A, U4C and G28A RNAs. The stretch of cytidine residues (nucleotides 41-44) that is the putative PCBP2 binding site is shown in green (insets). Three days post-electroporation of JFH-1_T_ RNA with the indicated mutations was electroporated into Huh-7.5 cells, **(B)** intracellular viral titers and (**C**) extracellular viral titers were assessed by FFU assay, and (**D)** viral RNA accumulation was assessed by RT-qPCR. (**E**) The proportion of packaged RNAs was ascertained by calculating the ratio of total FFUs **(B+C)** to viral genome copies **(D)**. The relative titers, RNA levels, and percentages of packaged virions in **(B-E)** were all normalized to the WT virus condition. (**F**) Relative intracellular and **(G**) extracellular viral titers, resulting from the infection of Huh-7.5 cells with mutant JFH-1_T_ viruses (MOI 0.05), two days post-transfection with siRNAs. The elative titers were all normalized to the control siRNA condition of each virus. Bars represent the mean of at least three independent replicates; error bars show the standard deviation of the mean. P-values were calculated with multiple paired t-tests (ns, not significant; * p < 0.05; ** p < 0.01; *** p < 0.001; **** p < 0.0001).

To this end, we made use of point mutations in the 5’ UTR that stabilize the SLII conformation, specifically U4C, G20A, and G28A (**Figure 5A** and **Figure S6**) (Chahal et al., 2019, 2021). The U4C and G20A mutations change the U-G wobble base pair at the base of SLI into a C-G (U4C) or U-A (G20A) Watson-Crick base pair, which results in similar energetic stabilities (ΔG) for the SLII^alt^ and SLII conformations. Meanwhile, the G28A mutation destabilizes SLII^alt^, rendering the SLII conformation comparatively more stable (Amador-Cañizares et al., 2018a; Schult et al., 2018; Chahal et al., 2019, 2021). To test if this stabilization of SLII altered virion assembly, we introduced each of the point mutations into the JFH-1_T_ genome and electroporated these RNAs into Huh-7.5 cells, then monitored intracellular and extracellular titers as well as HCV RNA accumulation (**Figure 5B-E**). Fascinatingly, we noticed an inverse correlation between the relative stability of SLII and intracellular and extracellular viral titers at day 3 post-electroporation (**Figure 5B-C**), yet we observed no significant differences in viral RNA accumulation (**Figure 5D**). Notably, the U4C and G28A viral RNAs consistently packaged a smaller proportion of their viral genomes that than the WT or G20A RNAs (**Figure 5E**). Consistent with this finding, and in agreement with our previous finding that PCBP2 knockdown only reduced the accumulation of viral RNAs that could engage in the early steps of virion assembly, PCBP2 knockdown had no effect on the accumulation of full-length G28A J6/JFH-1 RLuc reporter RNA (**Figure S8**). This is likely due to further impairments in virion assembly (in addition to the RLuc insertion) through stabilization of the SLII structure imposed by the G28A mutation. Moreover, we found that all the 5’ UTR mutant viruses (U4C, G20A and G28A) were similarly sensitive to PCBP2 knockdown during infection as WT (**Figure 5F-G**). Taken together, these results suggest that the SLII^alt^ conformation is an important requirement for efficient packaging of the genomic RNA into virions. Furthermore, while our findings suggest that SLII^alt^ and PCBP2 both modulate virion assembly, the SLII_alt_ conformation is either dominant or acts upstream of PCBP2 in the HCV assembly pathway, and thus modulates virion assembly independently from PCBP2.

## DISCUSSION

Herein, we investigated the role of PCBP2 in the HCV life cycle. We found that PCBP2 was not necessary for viral entry, translation, genome stability, RNA replication, or virion egress; but rather, endogenous PCBP2 only influenced the accumulation of viral RNAs that could complete early steps of the virion assembly process. Additionally, we found that mutations in the HCV 5’ UTR that destabilize SLII^alt^ conformations reduce the packaging efficiency of viral genomes into virions, independently of PCBP2’s role in the viral life cycle. Based on these findings, we propose a model whereby endogenous PCBP2 normally interferes with the transfer of the viral genomic RNA from NS5A to the core protein, thereby preventing premature virion assembly and indirectly promoting viral translation and RNA replication (**Figure 4G**). We further propose that the ability to form SLII^alt^ is required for efficient viral RNA packaging into virions, likely by preventing nascent viral genomes from engaging in translation (**Figure 6**).

**Figure 6.**
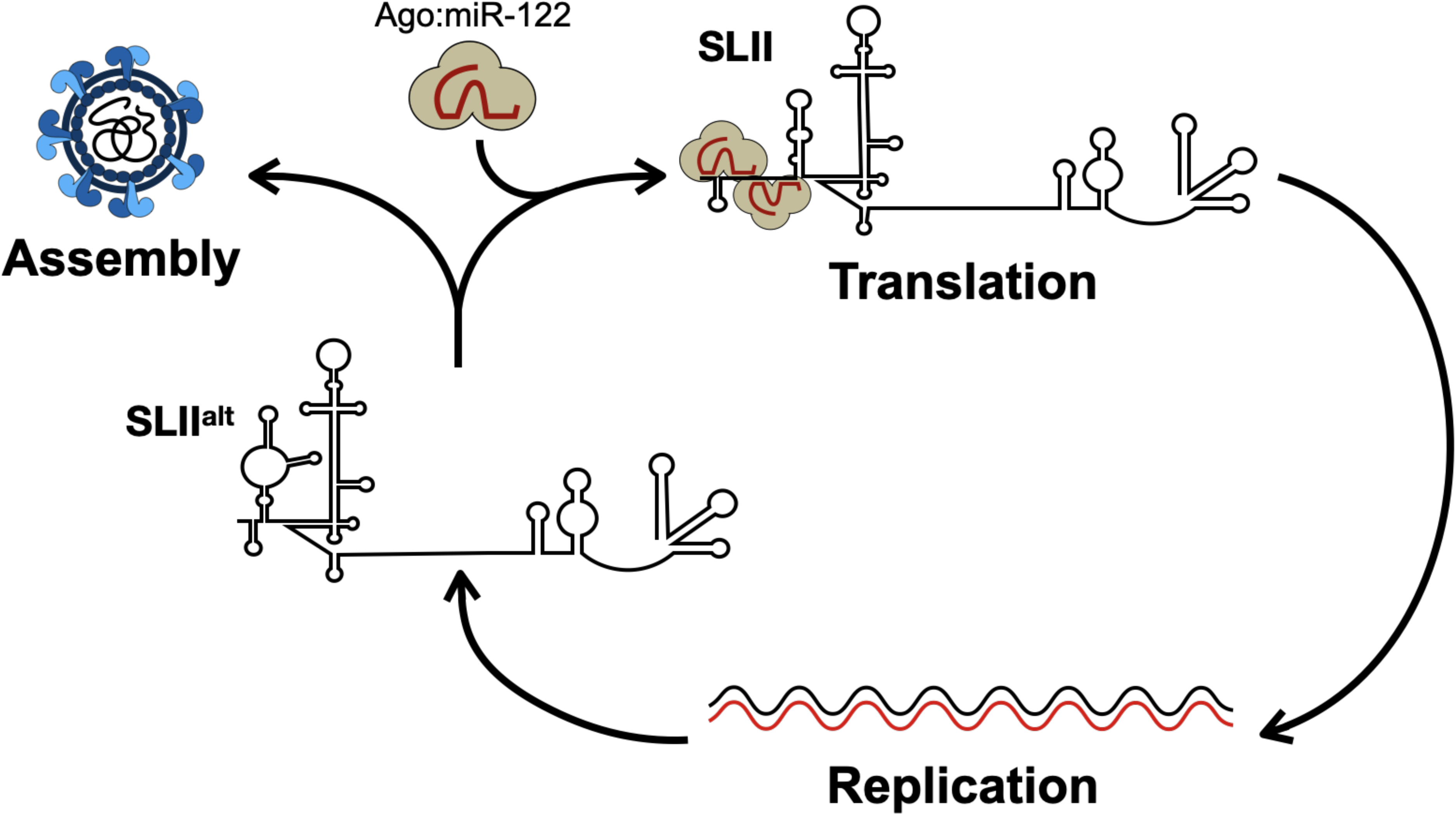
Model of the functional role of SLII^alt^ in the HCV life cycle. Newly-synthesized viral genomes are at a junction point in the HCV life cycle: they can either engage in virion assembly and be packaged into new virions that exit the cell, or they can engage in a new cycle of translation and RNA replication. Our model suggests that newly synthesized RNAs that take on the most energetically stable conformation, SLII^alt^, can thus engage in packaging. However, if they spontaneously or through binding of Ago:miR-122 convert to the functional SLII conformation, they form the viral IRES and recruit ribosomes, committing to translation.

Although HCV packaging and assembly are still incompletely understood, data to date suggest that it is a two-step process: 1) NS5A transfers the viral genome to the core protein at the surface of lipid droplets; and 2) a protein complex involving NS2, NS3, NS4A and the structural proteins (core, E1 and E2) simultaneously assemble the nucleocapsid and envelope it as it buds into the ER lumen (Lindenbach, 2013; Neufeldt et al., 2018; Masaki et al., 2008; Tellinghuisen et al., 2008; Appel et al., 2008; Jones et al., 2007, 2011; Roder et al., 2019). Based on our observation that PCBP2 knockdown only inhibits the accumulation of reporter RNAs with intact NS5A-core interactions, our data supports a model where PCBP2 limits virion assembly during the first step, whereby NS5A transfers the viral genomic RNA to core (**Figure 4G**). We favor this step rather than the overall process of virion assembly because ΔE1-p7 reporter RNAs were just as sensitive to PCBP2 knockdown as the WT full-length reporter RNA (**Figure 3A, C**). Since nucleocapsid formation occurs simultaneously with its envelopment, ΔE1-p7 viral RNAs cannot undergo appropriate packaging, but can still associate with the core protein, which is proposed to interact with numerous low affinity sites across the viral genomic RNA to mediate packaging (Stewart et al., 2016; Shi et al., 2016; Shimoike et al., 1999; Tanaka et al., 2000).

While the precise molecular mechanism by which PCBP2 interferes with the core-NS5A interaction remains unclear, based on previous studies and the data presented herein, we envision that PCBP2 could mediate its effects on the viral life cycle either by interacting with the viral proteins themselves or through direct interactions with the viral RNA. Firstly, as previous studies reported that PCBP2 binds to NS5A *in vitro* and can co-immunoprecipitate exogenously overexpressed NS5A, it is possible that PCBP2 interferes with viral RNA transfer through direct interactions with NS5A (Wang et al., 2011; Germain et al., 2014). PCBP2 interactions with NS5A may directly block NS5A interactions with the viral RNA or core protein, or could interfere with the NS5A phosphorylation events necessary for its delivery of the viral genome to core (Zayas et al., 2016). Alternatively, it is possible that PCBP2 interactions with the viral RNA may preclude NS5A or core from interacting with specific sites on the viral RNA. Notably, iCLIP analysis mapped a major PCBP2 binding site to the poly(U/UC) tract, a region of the genome known to be bound by NS5A; thus, it is possible that PCBP2 and NS5A may compete for binding to this region, or to other low affinity sites across the viral RNA important for packaging (Flynn et al., 2015; Foster et al., 2010; Stewart et al., 2016; Shi et al., 2016)

Importantly, our model does not require PCBP2 to play a direct role in translation or viral RNA replication to ultimately enhance intracellular viral protein expression or RNA accumulation. Our finding that PCBP2 has no direct effect on viral translation when assessed in isolation from the rest of the viral life cycle is consistent with prior studies that examined PCBP2’s effect on HCV IRES-mediated translation (Fukushi et al., 2001; Choi et al., 2004; Rosenfeld and Racaniello, 2005; Fontanes et al., 2009; Shirasaki et al., 2010). Furthermore, we found that PCBP2 was not necessary for viral RNA replication in the absence of virion assembly, which is consistent with a prior report that PCBP2 knockdown had no effect on the replication of a subgenomic HCV replicon that did not contain the *core* through *NS2* genes (Tingting et al., 2006). While our results are largely in agreement with prior studies, our findings are in contrast with those reported by Masaki *et al*., who found that silencing PCBP2 reduced nascent viral protein synthesis in a stably infected Huh-7.5 cell line (Masaki et al., 2015). However, a direct effect on protein synthesis is not necessarily supported by their data, which suggests that PCBP2 knockdown both impairs viral translation and decreases viral RNA accumulation to a similar extent, thereby reducing the overall quantity of templates available for translation (Masaki et al., 2015). Interestingly, their data suggested that PCBP2 knockdown did not reduce the rate of nascent viral RNA synthesis, suggesting that a similar number of viral replication complexes are present under PCBP2 knockdown and control conditions. This observation is compatible with the model proposed herein, as PCBP2 knockdown reduces the total intracellular RNA pool by promoting virion packaging, which is unlikely to disrupt the already established replication complexes in a persistently HCV-infected cell population.

There remains a lingering question as to whether PCBP2 and miR-122 compete or collaborate in their interactions with the HCV genome. Masaki *et al*. reported that miR-122 supplementation stimulates nascent viral RNA synthesis by competing with PCBP2 for the 5’ terminus of the viral RNA (Masaki et al., 2015). This competition has been more recently clarified by Gebert *et al*., who found that Ago:miR-122 complexes and PCBP2 specifically compete to bind to the miR-122 site 2 on the HCV genome, which overlaps with a stretch of four cytidines (C41-44) that appear to be the putative 5’ PCBP2 binding site (Gebert et al., 2021). However, it is important to note that these studies relied on a 47-nucleotide RNA probe of the HCV 5’ terminus, which is too short to form the SLII conformation and therefore leaves the C41-44 stretch single-stranded and thus accessible for PCBP2 binding. Given that the C41-44 stretch is only accessible for PCBP2 binding in the SLII^alt^ conformation(s), it is possible that PCBP2 and miR-122 only compete for binding to the viral RNA in this conformation, but not in the SLII conformation (where miR-122 interactions would predominate). However, additional binding studies using larger viral RNA fragments able to sample these different conformations are needed to further clarify this point. Notably, when we stabilized the SLII conformation with point mutations, we did not observe significant changes in PCBP2 sensitivity (**Figure 5F-G**). These findings suggest that either the RNA is still able to sufficiently sample the SLII^alt^ conformation(s), that PCBP2 can bind to the RNA in the SLII conformation, or that PCBP2’s activities are primarily regulated by its other interactions with the HCV genome, rather than this particular site at the 5’ terminus of the viral RNA.

Finally, in our attempt to clarify if the conformation of the viral RNA at the 5’ terminus was important for PCBP2 activity, our experiments revealed that the SLII^alt^ conformation is required for optimal virion assembly (**Figure 6**). This conclusion is based on our observation that mutations that stabilized SLII over SLII^alt^ reduced viral packaging efficiency in cell culture. More specifically, without significantly altering intracellular viral RNA accumulation, we found that the U4C and G28A mutations significantly reduced both intracellular and extracellular viral titers. The latter is consistent with a prior study, which reported that a G28A virus produced significantly lower infectious viral progeny than WT HCVcc, even in miR-122-replete conditions (Hopcraft et al., 2016). Herein, we also found that U4C, which stabilizes SLII without destabilizing SLII^alt^, reduced virion production but to a lesser extent than G28A (**Figure 5**).

Interestingly, the G20A mutation did not have any detectable impact on virion production, even though its secondary structure closely matches that of U4C (**Figure 5 and Figure S5**). Notably, both U4C and G20A stabilize the SLII conformation by replacing a G-U wobble with a canonical Watson-Crick base pair at the base of SLI: G20A generates an A-U pair, which has a similar stability to the G-U wobble, while U4C creates a G-C pair, which is thermodynamically more stable (Strazewski et al., 1999). Thus, it may be that although U4C or G20A are similarly likely to form the SLII^alt^ or SLII conformations, the structure of the G20A RNA may simply be more dynamic. Nonetheless, our results suggest a model whereby the SLII^alt^ conformation promotes virion assembly, while the SLII conformation promotes viral translation (**Figure 6**). However, since PCBP2 knockdown similarly affected each of these mutants, it appears that the SLII^alt^ conformation is either dominant or acts upstream of PCBP2 in the HCV assembly pathway, and thus modulates virion assembly independently from PCBP2.

Interestingly, our results may help explain the ability of the JFH-1 strain to produce infectious particles in cell culture (**Figure 7**). The JFH-1 strain was the first infectious HCV isolate to recapitulate the entire HCV life cycle in cell culture without adaptive mutations, including the production of infectious viral particles (Lindenbach et al., 2006; Wakita et al., 2005; Zhong et al., 2006). Subsequently, chimeric viruses that combined the JFH-1 (genotype 2a) genome with core-NS2 or 5’ UTR-NS2 sequences derived from genotypes 1-6 were developed and shown to produce infectious viral particles in cell culture (reviewed in (Wakita, 2019)). While the ability to produce infectious virions is certainly due at least in part to contributions from the JFH-1 nonstructural genes, our findings herein suggest that there may also be a contribution from the G residue at position 28 (G28) in the 5’ UTR. Notably, ∼80% of all HCV isolates contain an A at this position (A28) (Israelow et al., 2014). In JFH-1, the presence of G28 provides thermodynamic stability to SLII^alt^, and as we demonstrate herein, helps to prioritize virion assembly (**Figure 7A**) (Gottwein et al., 2009; Li et al., 2011; Pietschmann et al., 2006). In contrast, in other HCV genotypes, the A28 residue stabilizes the SLII conformation, which promotes viral translation to the detriment of virion assembly (**Figure 7B**). Moreover, the A28 residue also increases the affinity of the viral RNA for Ago:miR-122 at both miR-122 binding sites, which likely further reinforces a commitment to translation (Israelow et al., 2014; Chahal et al., 2021; Schirle and MacRae, 2012; Schirle et al., 2014; Sheu-Gruttadauria et al., 2019). Taken together, this suggests that while JFH-1 can prioritize virion assembly over translation, the other genotypes end up lost in translation. Furthermore, it is interesting to consider whether this may help explain HCV infection outcomes. JFH-1 was isolated from a patient with fulminant (acute) hepatitis, and several case reports have suggested an association between genotype 2 and fulminant hepatitis or severe recurrence post-transplant (Wakita et al., 2005; Kanzaki et al., 2014; Tracy et al., 2017). However, this association may be related to higher incidence of genotype 2 in these patient populations, as fulminant hepatitis cases have been reported with other genotypes (Sergi et al., 1998; Chu et al., 1999; Sakai et al., 2007). Moreover, despite its worldwide distribution, genotype 2 (G28) only accounts for an estimated 9.1% of HCV cases globally, and is highly responsive to interferon-free direct-acting antiviral regimens (Jacobson et al., 2013; Lawitz et al., 2013; Messina et al., 2015). It is unclear if the lower prevalence of genotype 2 is related to a higher rate of spontaneous viral clearance or to other factors as genotype 2 has been sporadically reported to be less likely to progress to chronic infection — with estimates between 22-33% for genotype 2 versus 57-92% for genotype 1 (Amoroso et al., 1998; Cho et al., 2014). However, spontaneous clearance rates between genotypes are unreliable, as they largely rely on the assessment of outbreak studies of single genotypes in distinct patient populations and are thus subject to cohort bias (Amoroso et al., 1998; Cho et al., 2014; Messina et al., 2015). Nonetheless, it is possible that more efficient replication and infectious particle production generates a more robust intracellular and systemic immune response that can effectively control HCV infection at the acute stage. In contrast, prioritizing translation to the detriment of virion assembly may allow HCV to better evade intracellular antiviral responses and avoid induction of a systemic immune response, thereby favoring the establishment of a chronic infection. Consistent with this idea, treatment-naïve chronic genotype 2 patients have been demonstrated to have greater HCV-specific T cell responses at baseline, which could be reflective of early viral infection kinetics (Kaplan et al., 2005). However, more research is needed to provide further insight into whether G28 status influences disease pathogenesis.

**Figure 7.**
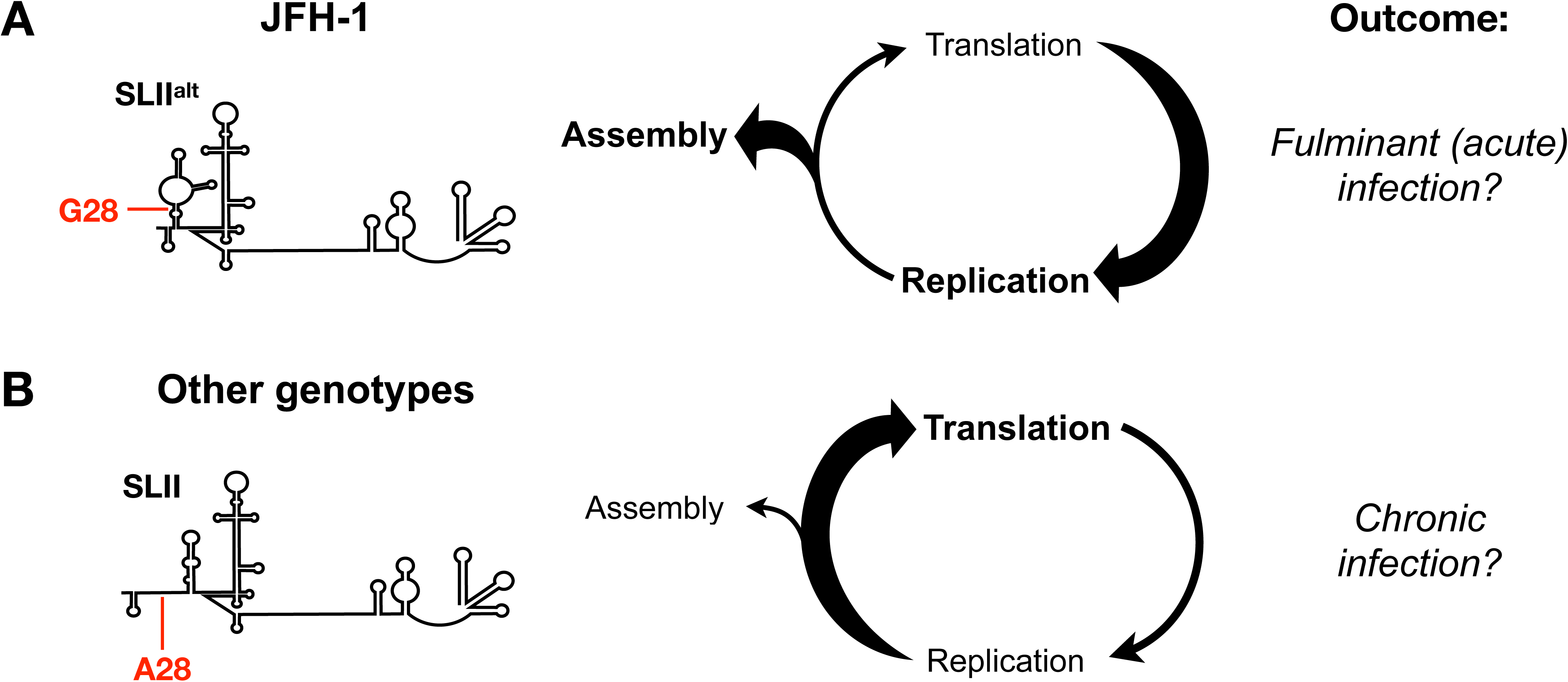
The formation of SLII^alt^ may explain why JFH-1 is able to produce virions in cell culture and has implications for the outcome of HCV infections. **(A)** Previous studies indicate that JFH-1 non-structural proteins provide it with a replicative advantage over other HCV isolates. Additionally, the G28 residue in the 5’ UTR allows JFH-1 (and likely other genotype 2 isolates) to energetically favor the SLII^alt^ conformation, which we demonstrate promotes virion assembly. This prioritization of virion assembly over translation may contribute to JFH-1 pathogenesis, considering it was isolated from a patient with fulminant (acute) hepatitis and genotype 2 has been associated with increased incidence of fulminant hepatitis. **(B)** In contrast, the other HCV genotypes (∼80% of all HCV isolates) have an A at position 28 (A28), which energetically favors the SLII conformation rather than SLII^alt^. As a result, these other genotypes favor translation over virion assembly. Moreover, A28 increases the affinity of the 5’UTR for Ago:miR-122 complexes, further reinforcing translation. This bias may allow the virus to better evade intracellular and systemic immune responses, thereby resulting in chronic HCV infection.

In conclusion, our results clarify the role of PCBP2 in the HCV life cycle and support a model where endogenous PCBP2 inhibits the early steps of virion assembly mediated by the core and NS5A proteins. By preventing viral genome sequestration by the core protein, PCBP2 indirectly promotes viral RNA retention in the translating/replicating pool, which may explain previously reported discrepancies with respect to PCBP2’s effects on HCV translation and viral RNA replication. Furthermore, we found that the ability of the viral genomic RNA to form the SLII^alt^ conformation was important for HCV RNA packaging, in a manner that is independent of PCBP2s role in virion assembly. Our data suggests that the ability to form SLII^alt^ may partially explain infectious particle production in HCVcc models and may offer insight into the outcome of HCV infection. Taken together, our results provide models for how SLII^alt^ and PCBP2 independently modulate HCV genome packaging and alter the balance between viral RNAs in the translating/replicating pool and those engaged in virion assembly.

## ACKNOWLEDGEMENTS

We would like to acknowledge Charlie Rice (Rockefeller University) for kindly providing the Huh-7.5 cells, pJ6/JFH FL RLuc WT, and GNN plasmids; Rodney Russell (Memorial University) for providing JFH-1_T_; Mamata Panigrahi and Joyce Wilson (University of Saskatchewan) for the pJ6/JFH mono RLuc NS2 and pJ6/JFH-1 E1-p7 del plasmids; Martin J Richer (McGill University) for the 293T cells; and John Law (University of Alberta) for the kind gift of HCVpp. We are also grateful to Nathan Taylor, Julie Magnus, Carolina Camargo and Marylin Rheault (McGill University) for technical support. Finally, we are grateful to Bert Semler and Hung Nguyen for their insights into PCBP2-RNA interactions, and to Sonya MacParland and Jordan Feld for critical discussions regarding clinical outcomes of viral infection.

## AUTHOR CONTRIBUTIONS

S.E.C. and S.M.S. designed the study; S.E.C. performed the experiments and analyzed the data, and S.E.C and S.M.S. wrote and edited the manuscript.

## SUPPLEMENTARY INFORMATION SUPPLEMENTARY METHODS

### IRES-specific translation assays

#### Cells

HeLa cervical epithelial adenocarcinoma cells obtained from the ATCC (CCL-2) were maintained in DMEM supplemented with 10% FBS, at 37°C/5% CO_2_, and were routinely screened for mycoplasma contamination.

#### Plasmids

Plasmids encoding a firefly luciferse (FLuc) reporter gene under the translational control of a poliovirus IRES (PV IRES, nt 71-732), HCV IRES (nt 40-372) or encephalomyocarditis virus IRES (EMCV IRES, nt 281-848) were a kind gift from Drs. Yuri Svitkin and Nahum Sonenberg (Svitkin et al., 2001). These plasmid templates were linearized with *BamHI* and *in vitro* transcribed using T7 RNA polymerase as previously described (Amador-Cañizares et al., 2018b). The Renilla luciferase (RLuc) mRNA was transcribed from the pRL-TK plasmid (Promega) linearized using *BglII* and *in vitro* transcribed using the mMessage mMachine T7 Kit (Life Technologies) according to the manufacturer’s instructions.

#### Transfection and luciferase assay

HeLa cells were plated at a density of 5 x 10^5^ cells/10-cm dish and transfected with 50 nM of siPCBP2 or siCTRL (miR-122_p2-8_) siRNA duplexes, as described in the main text’s methods. The next day, transfected cells were seeded into 6-well plates at a density of 2.5 x 10^5^ cells per well. On the second day post-siRNA transfection, each well was co-transfected with 1.5 µg of IRES-FLuc IVT and 2.5 µg of RLuc mRNA, resuspended into 1 mL OptiMEM with 2.5 µL of Lipofectamine 3000 (Invitrogen). Four hours post-transfection, cell culture media was changed for DMEM with 10% FBS. The next day, each well was harvested in 100 µL of 1X passive lysis buffer (Promega). Renilla and firefly luciferase activity was measured using the Dual-Luciferase Assay Reporter Kit (Promega), following the manufacturer’s instructions with the modification that 25 µL of reagent were used with 10 µL of sample. All samples were measured in triplicate.

### Subgenomic replicon and RLuc-G28A plasmids

The pSGR FLuc WT plasmid bears a subgenomic replicon derived from JFH-1, where the core through NS2 region has been replaced by a firefly luciferase (FLuc) reporter gene and with an EMCV IRES, which drives the translation of the NS3 through NS5B proteins (Wakita et al., 2005); this plasmid was provided by Dr. Ralf Bartenschlager. The J6/JFH RLuc G28A plasmid, which bears a full-length J6/JFH RNA with the RLuc reporter gene and with a G28A mutation in its 5’ UTR, had been previously cloned by Dr. Jasmin Chahal (Chahal et al., 2021). Both plasmids were linearized by *XbaI* for *in vitro* transcription with T7 RNA polymerase.

## SUPPLEMENTARY FIGURES

**Figure S1.**
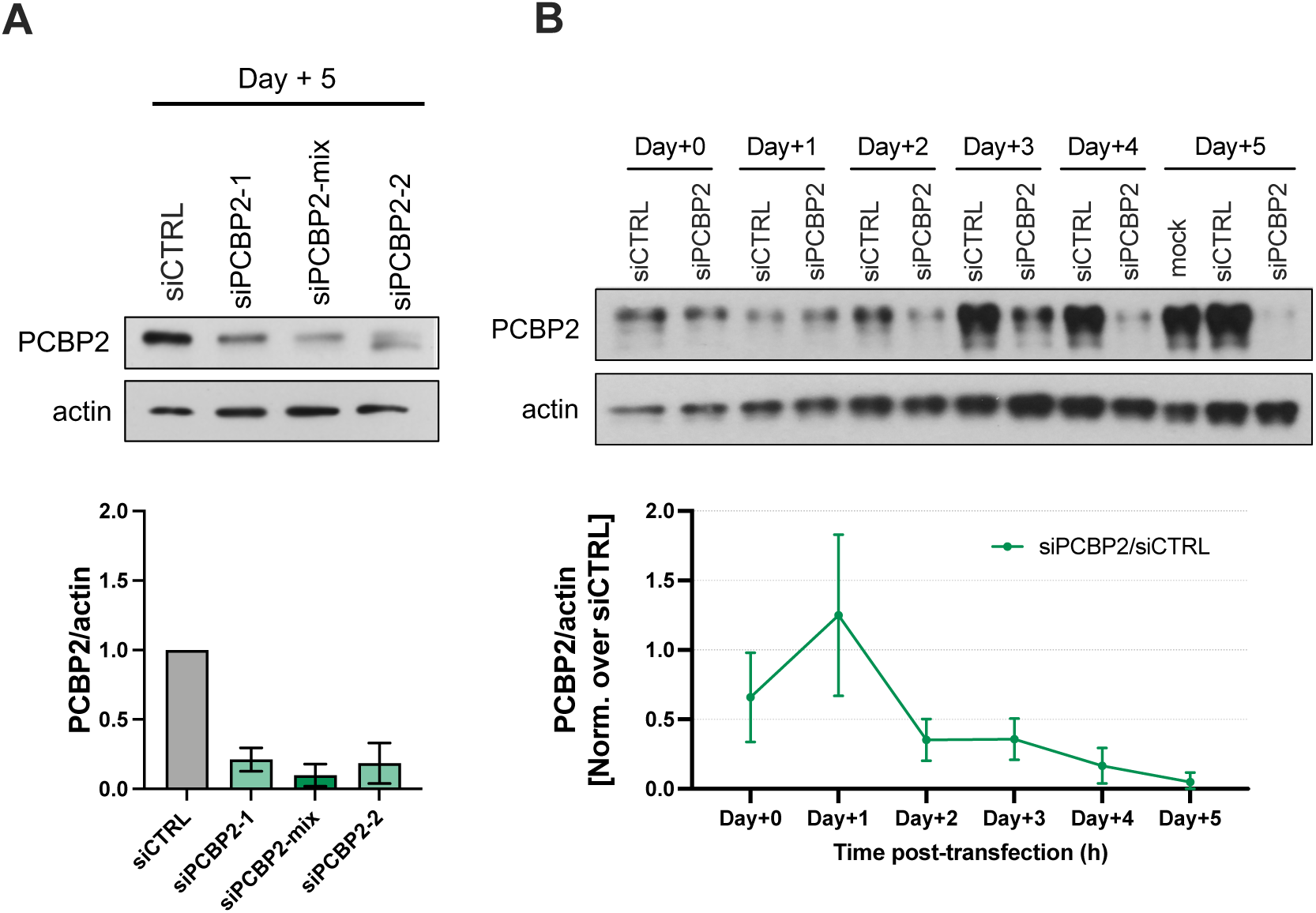
Optimization of the anti-PCBP2 siRNA transfection. **(A)** Comparison between single-siRNA transfections and the 1:1 combination of both siRNAs (siPCBP2-mix). The error bars (below) represent the standard deviation from two independent experiments. **(B)** Analysis of PCBP2 knockdown over time, compared to the siCTRL condition. PCBP2/actin band densities were compared to the same day’s siCTRL condition (line graph, below); error bars represent the standard deviation from two independent replicates.

**Figure S2.**
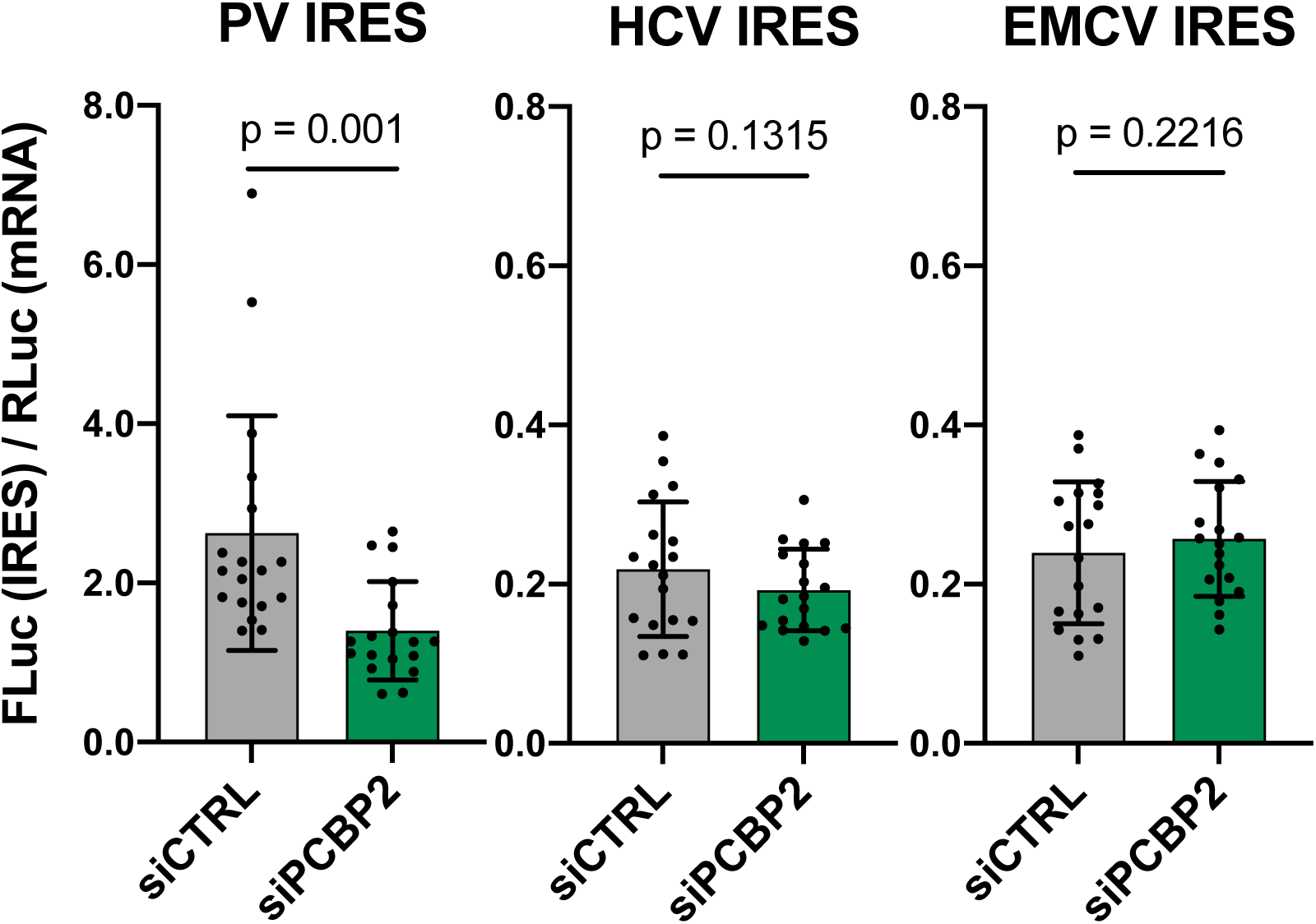
PCBP2 knockdown decreases PV IRES-mediated translation, but has no effect on HCV or EMCV IRES-mediated translation. Two days post-siRNA transfection, HeLa cells were co-transfected with 2.5 µg of capped Renilla luciferase mRNA and 1.5 µg of RNAs comprised of a firefly luciferase (FLuc) gene under the control of either the PCBP2-sensitive poliovirus (PV) IRES, the PCBP2-insensitive encephalomyocarditis virus (EMCV) IRES, or the hepatitis C virus (HCV) IRES. Total protein samples were harvested in passive lysis buffer one day later, and luciferase activity was determined using the Dual Luciferase Reporter Assay kit. The IRES-mediated translation signal (FLuc) was normalized to the transfection efficiency control (RLuc) signal. Bars represent the mean ± standard deviation of four independent replicates. P-values were determined by paired t-test.

**Figure S3.**
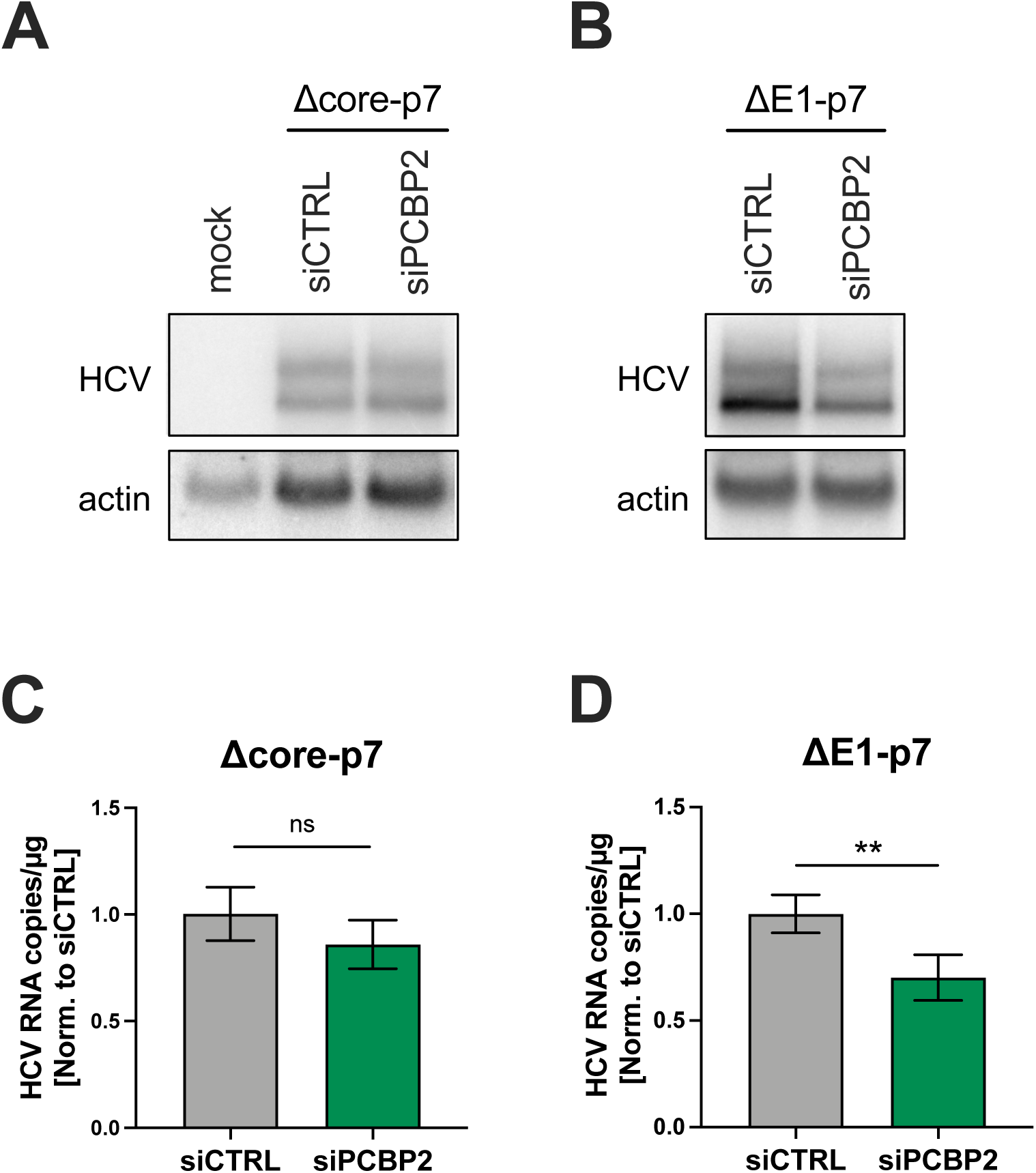
PCBP2 knockdown does not affect Δcore-p7 RNA accumulation, but decreases ΔE1-p7 RNA accumulation. **(A)** Two days post-electroporation into Huh-7.5 cells, Δcore-p7 and **(B)** ΔE1-p7 RNA accumulation was assessed by Northern blot. **(C)** Δcore-p7 RNA accumulation and **(D)** ΔE1-p7 RNA accumulation was also assessed by RT-qPCR. Error bars represent the standard deviation of two independent Δcore-p7 replicates or three independent ΔE1-p7 replicates. P-values were determined by paired t-test (ns, not significant; ** p < 0.01).

**Figure S4.**
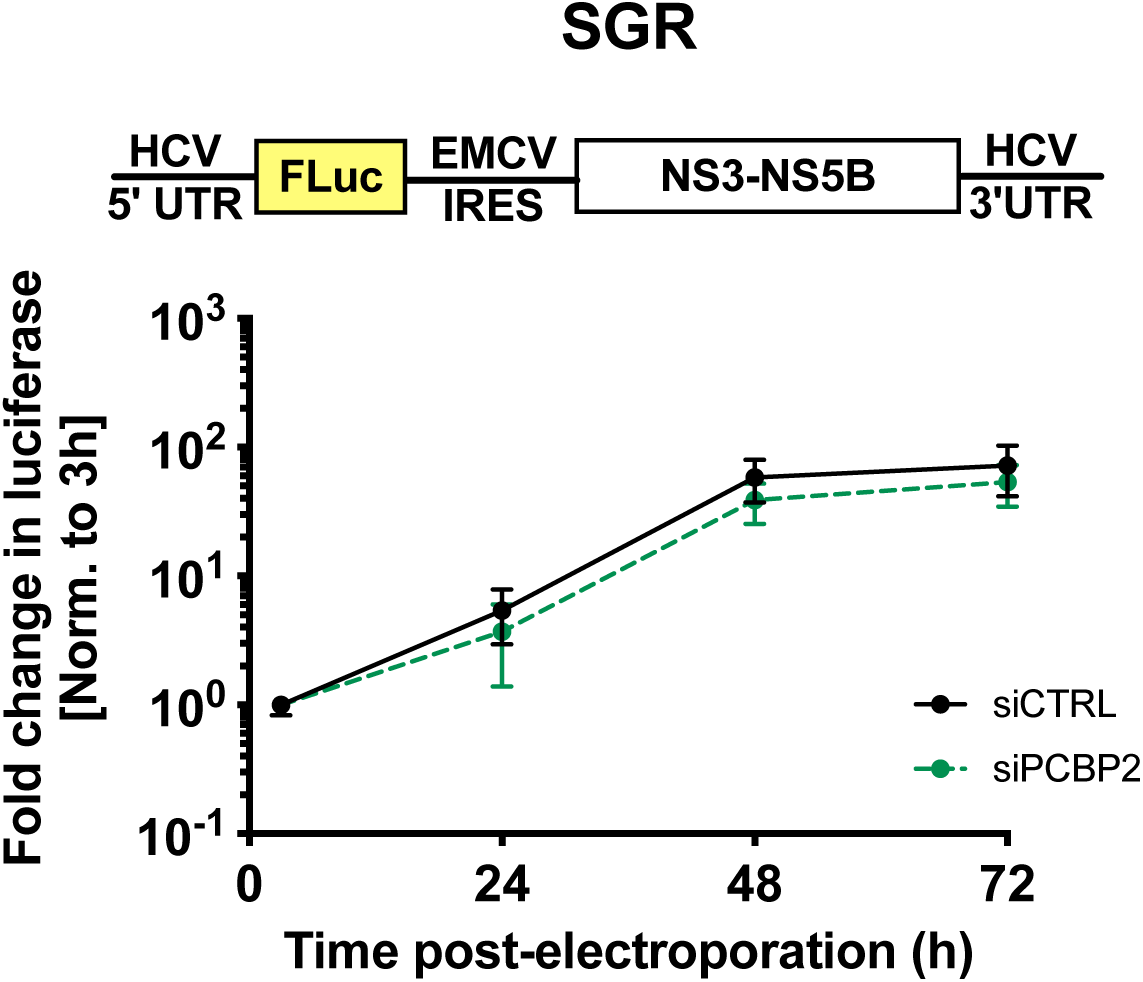
PCBP2 knockdown does not affect the replication of the bicistronic subgenomic replicon. Two days post-siRNA transfection, Huh-7.5 cells were electroporated with 10 µg of reporter bicistronic subgenomic replicon (SGR), and reporter luciferase gene activity was tracked for three days post-electroporation. FLuc values were normalized to the early timepoint (3 h), to control for disparities in electroporation efficiency between experiments. Data are representative of three independent replicates; error bars represent the standard deviation of the mean. P-values were calculated by two-way ANOVA.

**Figure S5.**
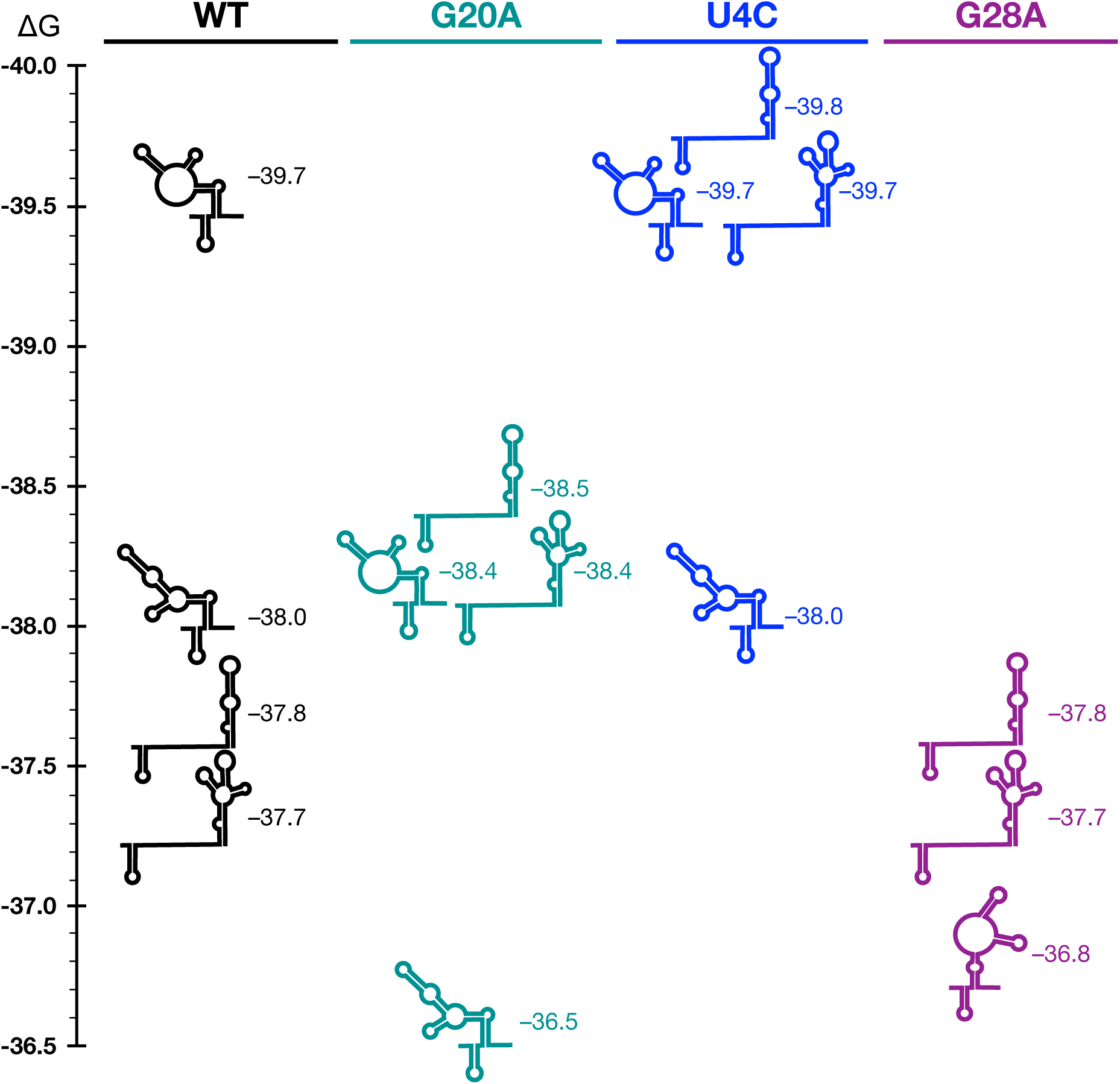
RNA structure predictions for the first 117 nucleotides of the JFH-1 genome, for WT, G20A, U4C and G28A sequences. RNA structure and their associated free energy (ΔG) predictions were calculated using RNAStructure, with the first 3 nucleotides constrained as single-stranded.

**Figure S6.**
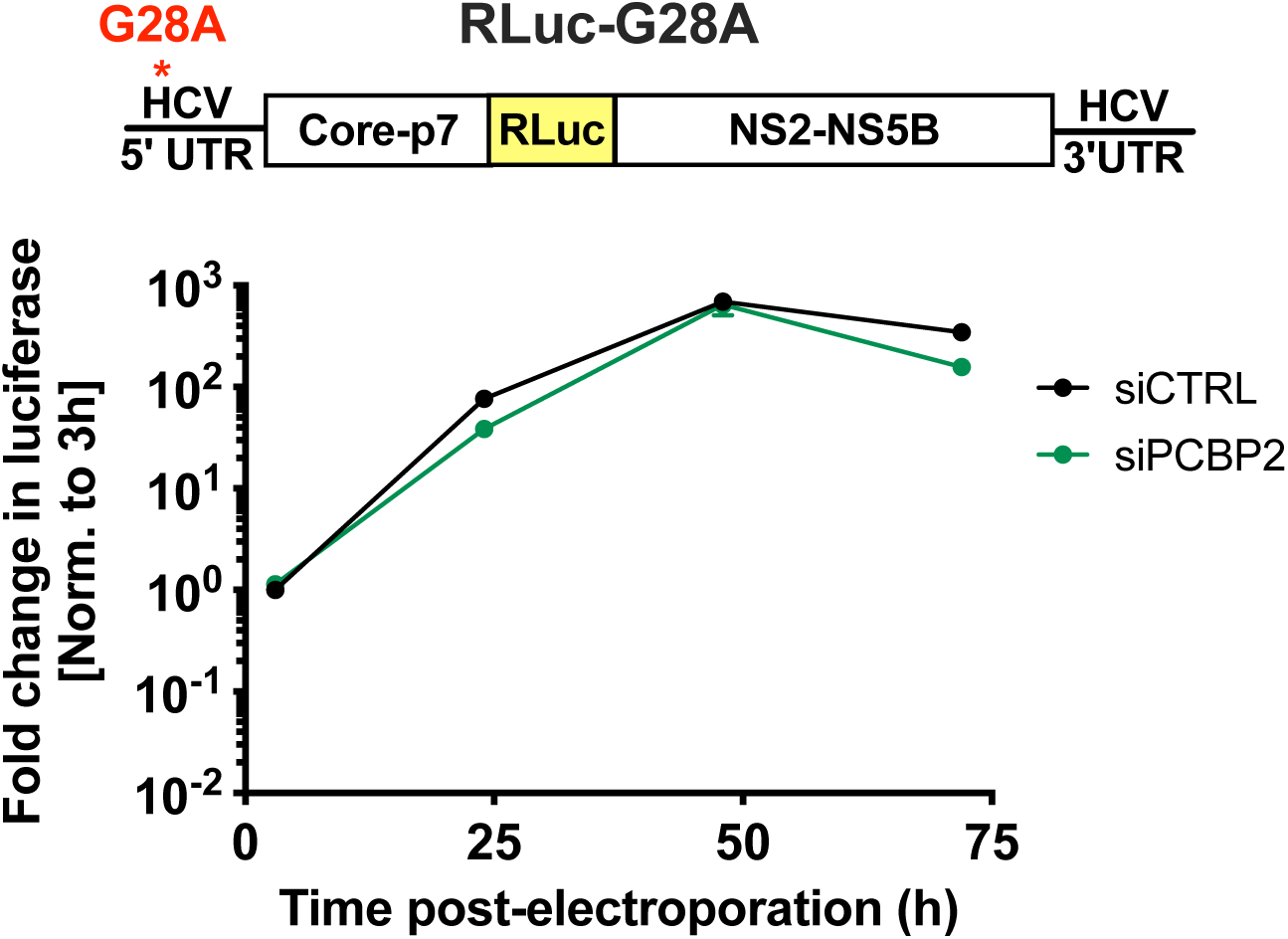
PCBP2 knockdown does not impact replication of a G28A mutant. Two days post-siRNA transfection, the full-length G28A J6/JFH-1-RLuc RNA (which is identical to the full-length J6/JFH-RLuc WT RNA shown in Figure 3A, but with a G28A substitution) was electroporated into Huh-7.5 cells, and luciferase activity was monitored at several timepoints post-electroporation. Data are representative of three independent replicates, and error bars represent the standard deviation of the mean.

